# mRNAfold: Co-optimization of Global Stability, Local Structure, and Codon Choice via Suboptimal Folding

**DOI:** 10.64898/2026.01.23.701221

**Authors:** Max Ward, Mary Richardson, Haining Lin, Michael Stamm, Kathryn Wright, Angie Kim, Alicia Bicknell, Nabeel Ahmed, Adriana Jones, J. Wade Davis, Mihir Metkar

## Abstract

mRNA medicines hold great promise, but designing sequences with high translation efficiency, robust in-solution stability, and manufacturability remains a major challenge due to the vast combinatorial space of synonymous coding sequences. Computational approaches such as mRNA folding algorithms have emerged as powerful tools by co-optimizing for in-solution stability and translation efficiency, yet current methods face important limitations. Here, we present “`mRNAfold`”, an improved mRNA folding algorithm and software package that addresses these gaps by enabling efficient exploration of diverse near-optimal solutions, incorporating untranslated regions (UTRs), parallel execution, and supporting tunable control over local structural features across the mRNA. Thermodynamically optimized mRNAs from `mRNAfold` were more stable (≈ 2-fold) in-solution than those generated by simple GC maximization for the same encoded protein. In addition, mRNAs designed to vary local structure near the start codon while maintaining consistent structure and codon optimality elsewhere showed a complex relationship between local structure near the start codon and protein production in cells. We observed no impact of structure in the start codon region for a set of mRNAs with high codon optimality, but it did impact protein production for a set of mRNAs with lower codon optimality. Together, these results underscore the potential of structure-aware, multi-objective design to improve mRNA medicines and offer a framework for exploring how sequence, structure, and expression are interrelated.

## 1 Introduction

Recent advances in mRNA medicines have been transformative. The advent of mRNA vaccines is reshaping the landscape of infectious disease control [9, 13]. Moreover, mRNA’s capacity to encode customizable proteins holds promise for applications in cancer therapy, personalized medicine, genome editing, and protein replacement [34]. For commercial use, an ideal mRNA must be scalable to manufacture, stable during synthesis, storage, and distribution, and capable of efficient translation to produce sufficient protein in target cells. Remarkably, many of these properties can be engineered directly through the mRNA’s primary sequence [25, 28]. The central challenge, therefore, lies in designing sequences that effectively balance and encode all of these desirable features.

Current mRNA sequence design strategies often prioritize codon optimization to enhance protein expression. This is typically achieved by selecting synonymous codons that either match host tRNA abundance (tRNA adaptation index; tAI) [29] or mirror codon usage in mRNAs reference genes (Codon Adaptation Index; CAI) [33]. However, this approach neglects the RNA structural context, which is increasingly recognized as a key regulator of translation initiation, elongation dynamics, ribosome pausing, and transcript degradation [2, 3].

In addition to influencing cellular metabolism, RNA structure also impacts the stability of mRNA drugs in storage. Structured RNAs degrade more slowly in solution due to lower exposure of phosphodiester linkages to hydrolysis, an important consideration in the real-world deployment of mRNA vaccines [14]. During the COVID-19 pandemic, ultra-cold chain requirements posed considerable logistical barriers, especially in low-resource settings [39]. Thus, rationally designing mRNAs with more stable structures, achieved through increasing base-pairing, offers a promising strategy to improve shelf-life and reduce dependence on cold-chain logistics [42, 25].

However, designing sequences with optimized structure is computationally challenging. The number of synonymous mRNA sequences encoding a single protein grows exponentially with protein length, resulting in a vast combinatorial space. While codon optimality can be computed easily on a per-codon basis, RNA structure prediction is more complex. On average, a protein sequence has roughly 3^*M*^ synonymous mRNA variants (assuming 3 synonymous codons per amino acid on average), each of which can adopt approximately 2.6^*N*^ possible secondary structures [36], where *M* is the length of a protein in amino acids and *N* is the nucleotide length of the mRNA. To put this in context, a small protein sequence with *M* = 50 and *N* = 150, has ≈ 3^50^ × 2.6^150^ ≈ 10^86^ combinations. This is roughly the same magnitude as the number of elementary particles in the observable universe. For protein sequences used in therapeutic applications, the numbers are unfathomably larger.

To meet these challenges, a new class of mRNA design algorithms has emerged, aiming to co-optimize for both high CAI and high structural stability. An important subclass of these algorithms, termed mRNA folding algorithms, includes LinearDesign [45], DERNA [12], CDSFold [37], and the Cohen-Skiena algorithm [4], all of which extend dynamic programming formulations originally developed for RNA secondary structure prediction [47].

In our recent review of mRNA folding algorithms [41], we concluded that they offer a promising framework for rational mRNA design. These methods scale to long sequences, support joint optimization of codon usage using CAI and structural stability using Minimum Free Energy (MFE), and build on well-established RNA structure models. However, we also identified several key limitations that restrict their broader applicability in real-world design scenarios.

First, existing algorithms do not support diverse suboptimal folding, a core feature of classical RNA folding tools [44, 7]. Suboptimal folding methods are essential for navigating the uncertainty inherent in structure prediction and for exploring multiple design solutions. They have been around almost as long as RNA structure prediction algorithms have [46, 35]. In contrast, mRNA folding algorithms typically only output a single solution per run. The exception is DERNA, which outputs a narrow set of Pareto-optimal solutions, often yielding limited diversity. For example, Gu et al. report a median of only 18 solutions per protein sequence [12], making it difficult to obtain diverse sequences with both high CAI and low MFE.

Second, no current mRNA folding algorithm supports fully controlled local structure tuning. The ability to systematically guide structure formation in specific regions remains limited and is largely based on trial-and-error heuristics. Excessive secondary structure within the 5’UTR or at the start of the coding sequence (CDS) can inhibit ribosomal scanning, initiation efficiency, and start codon recognition [40]. Increased secondary structure within the CDS correlates with enhanced mRNA half-life in cells, resulting in greater protein output over time [22, 2]. Thus, being able to modulate structure locally could offer finer control over translation efficiency, fidelity, and duration. Existing approaches, like the LinearDesign heuristic that excludes the 5’ leader region from optimization and reintroduces it post hoc [45], are limited in scope and not well-integrated into available software.

Third, no current mRNA folding algorithm is designed to run in parallel. While the search space grows with sequence length, especially for real-world therapeutic designs, existing tools remain limited to single-threaded implementations.

Fourth, current methods exclude untranslated regions (UTRs), even though these are critical to the overall structure and function of the mRNA molecule [16, 23]. Optimizing only the coding sequence can lead to incomplete or even misleading design results, particularly when structural elements span CDS-UTR boundaries.

In this work, we present a new algorithmic toolkit “`mRNAfold`” that addresses all of the above limitations. Our toolkit supports a novel suboptimal folding algorithm, including a stochastic method that gives fine-grained control over sample diversity. The new mRNA folding algorithm we introduce is designed to run in parallel making use of multiple threads. Our toolkit is faster than existing algorithms, and much faster with multiple threads. It supports both MFE and CAI optimization, incorporates UTRs during optimization, and also includes an effective method to find optimized sequences with high overall structure, but decreased structure in target unpaired regions. Together, these capabilities enable the design of synthetic sequences to test a broader range of established and emerging hypotheses in mRNA design.

## 2 The mRNAfold Algorithm

### 2.1 Design Objectives

The goal of our algorithm is to design a valid CDS that balances RNA structural stability and codon usage for the target protein. Existing mRNA folding algorithms use MFE as a proxy for in-solution stability [37, 45, 12, 4, 42]. Lower MFE corresponds to high thermodynamic stability, which corresponds to higher in-solution stability. In addition, modern mRNA folding algorithms incorporate CAI [33] as a measure of codon optimality, a known predictor of protein expression [45, 12]. Following the approach employed by recent mRNA folding algorithms, we define an objective score that combines MFE and CAI. Let *π* be an RNA sequence. We use the following combined objective score originally from [45] with this version derived in [41]:

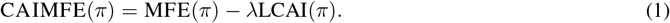

The relevant definitions are given in Table 1. A detailed derivation of these terms can be found in the Supplementary Materials.

**Table 1:**
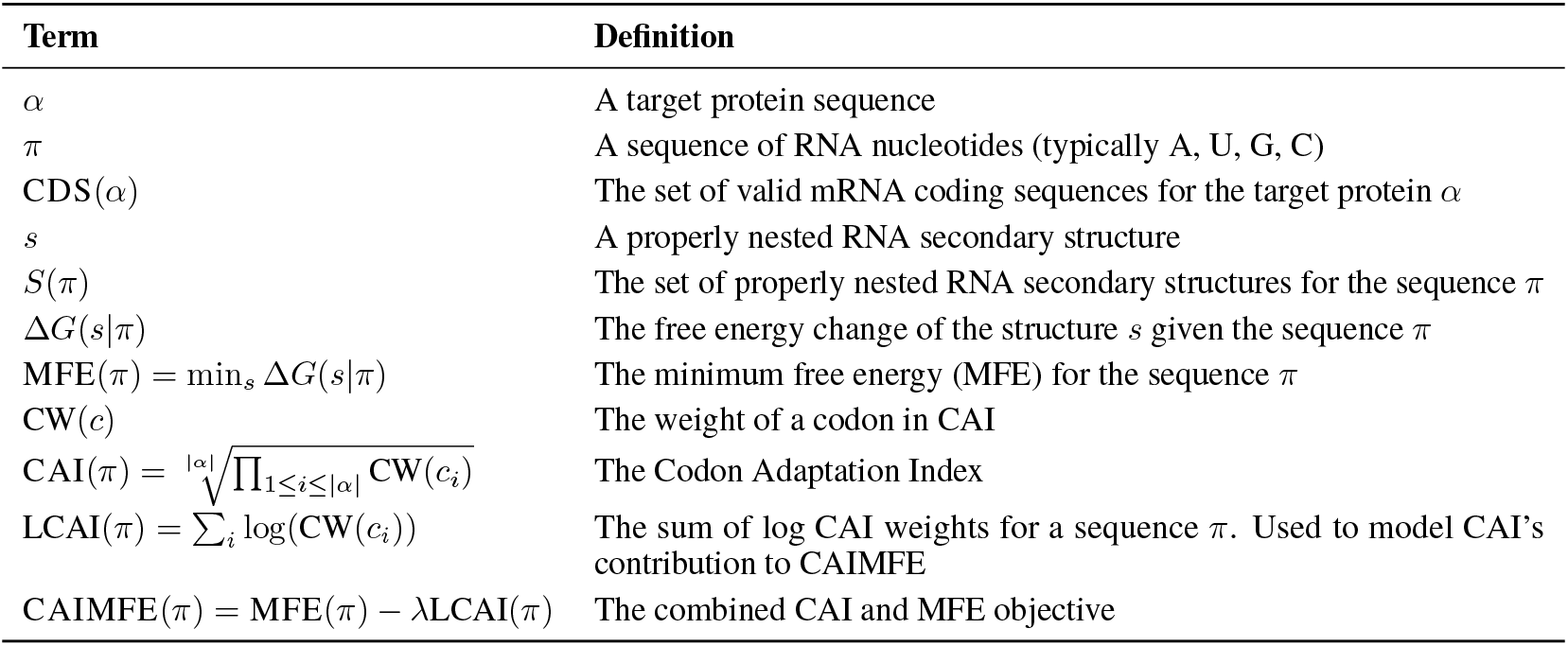
Definitions for mRNA Folding.

The goal of our mRNA folding algorithm is to minimize CAIMFE, resulting in a sequence with high stability (low MFE) and high codon optimality (high CAI). Formally, for a given protein sequence *α* with a set of valid CDSs CAI(*α*), we seek the RNA sequence *π* ∈ CAI(*α*) that optimizes CAIMFE:

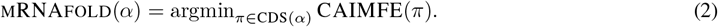

### 2.2 Nearest Neighbor Model Details

Our algorithm, like existing mRNA folding algorithms, extends the Zuker-Stiegler dynamic programming recurrences for RNA structure prediction [47]. These recursions, along with the Nearest Neighbor (NN) model, enable calculation of the MFE secondary structure for a given RNA sequence [38, 21, 20, 26].

We adapt the notation of our previous work [41] to describe the energy change functions in the NN model (Table 2). The energy of an RNA secondary structure is computed by summing the energetic contributions of its constituent loops, as defined by the NN model. These are classified into three categories: one-loops, closed by a single base pair; two-loops, which are closed by two base pairs; and multiloops, which are closed by more than two base pairs. The energy of one-loops and two-loops is dependent on the specific identities of the nucleotides forming the closing base pairs (e.g., A, U, G, C). The energy of a multiloop is treated with a linear model, comprising a constant initiation penalty plus terms that scale with the number of unpaired nucleotides and the number of closing pairs it contains. For conceptual clarity, this description omits advanced energetic terms such as coaxial stacking, loop-size initiation terms, dangling ends, end-penalties, terminal mismatches, and special loops with unique energies [26, 38]. We believe these are straightforward to incorporate once the algorithmic ideas are understood and are omitted only for clarity. We have also discussed the inclusion of some of these terms in the provided energy functions previously [41].

**Table 2:** Free energy change terms in the simplified Nearest Neighbor model we adopt.

| Loop Type | Notation | Energy Contribution |
| --- | --- | --- |
| One-loop | $\text{ONELOOP}(b_i, b_j)$ | The free energy contribution of a loop closed by one base pair (i.e., a hairpin loop). Dependent on the base identities of the closing pair $(b_i, b_j)$ |
| Two-loop | $\text{TWOLOOP}(b_i, b_j, b_k, b_l)$ | The free energy contribution of a loop closed by two base pairs (i.e., a stack, bulge, or internal loop). Dependent on the base identities of the two closing pairs $(b_i, b_j)$ and $(b_k, b_l)$ |
| Multiloop | $\text{ML}_{\text{init}} + \text{ML}_u \times u + \text{ML}_c \times c$ | Multiloop energy is the sum of an initiation term $\text{ML}_{\text{init}}$ plus a constant factor $\text{ML}_u$ for each unpaired nucleotide in the multiloop plus a constant factor $\text{ML}_c$ for each base pair closing the multiloop. Dependent on the number of unpaired nucleotides ($u$) and the number of closing pairs ($c$) in the multiloop. |

### 2.3 Codon Graphs

A key difference between mRNA folding algorithms, like our new algorithm, and RNA folding algorithms is that mRNA folding considers many possible sequences. RNA folding algorithms assume a fixed RNA sequence and operate on indexes into this sequence. Our algorithm consider all valid CDSs for a given protein. To achieve this, our algorithm uses pointers to nodes in a “codon graph”, similar to existing mRNA folding algorithms (Figure 1A). Each path through the codon graph corresponds to a valid CDS.

**Figure 1.**
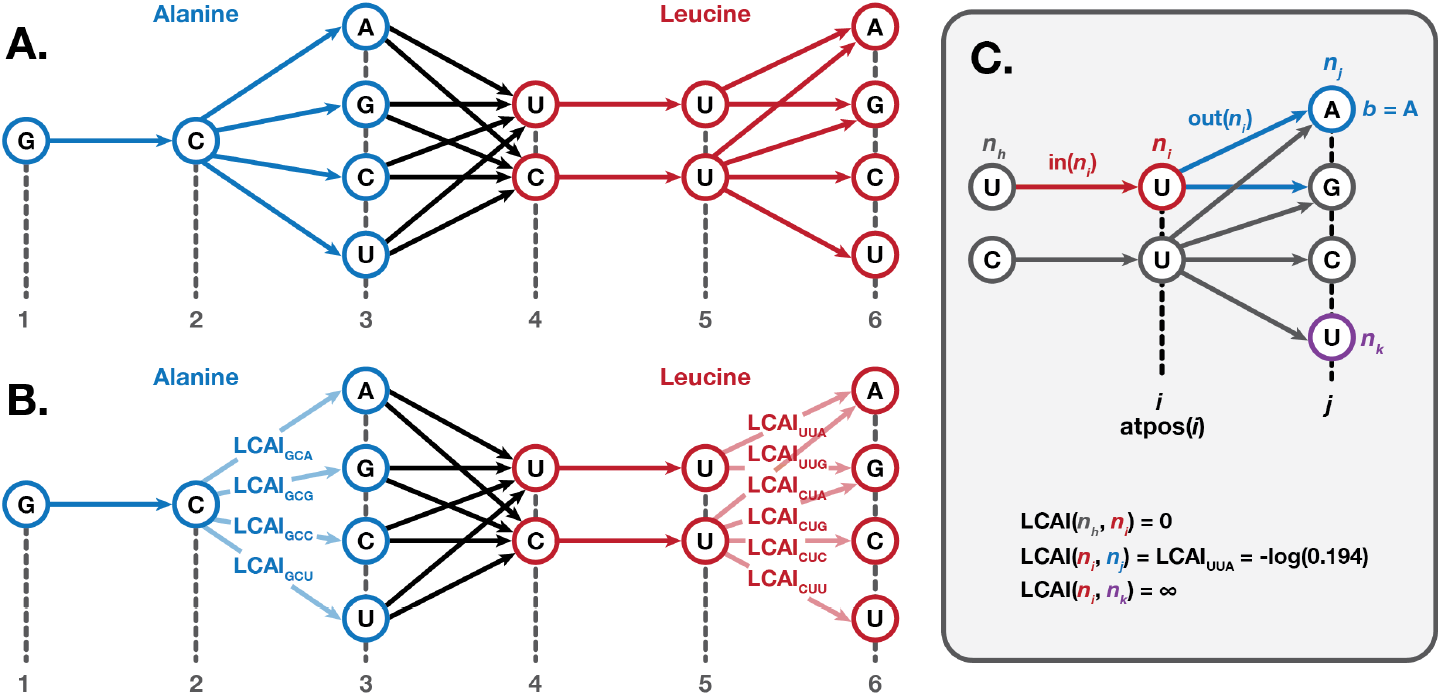
Codon graphs. (A) A codon graph considers all possible CDSs for the target protein. Each path through the graph corresponds to a single valid codon sequence. (B) Negative log-CAI weights (LCAI) are added on the third edge of each codon in the graph. (C) Each nucleotide in the codon graph can be accessed using pointers to specific nodes. Terms for accessing parts of the leucine subgraph are shown. Note that leucine (along with serine and arginine) represents the most complex case of converting the codon table to a codon graph, since there are six possible codons for this amino acid.

Note that both CDSfold [37] and LinearDesign [45] use similar codon-graph-like methods. CDSfold uses “extended nucleotides” to capture codon constraints, while LinearDesign uses an automaton. Neither is an explicit codon graph, but our review [41] points out that the two approaches are equivalent in practice and can be unified under a codon-graph framework.

#### 2.3.1 Incorporating CAI

Our algorithm incorporates CAI using a codon graph equivalent to LinearDesign’s automaton [41, 45]. This graph construction gives the rightmost edges for a codon a non-zero weight based on their log(CW(*c*)) value. Since RNA folding algorithms conventionally compute a minimum rather than a maximum, the negative log-CAI weights are used in the codon graph (Figure 1B).

Formally, define LCAI(*u, v*) as the negative log-CAI weight of the edge from node *u* to node *v*: LCAI(*u, v*) = − log (CW(*c*)). If no edge exists, then LCAI(*u, v*) = ∞. If the edge exists but does not have an associated weight, LCAI(*u, v*) = 0. In our codon graph, weights are associated with the last edge for each codon. However, this is not a strict constraint–weights could be associated with other edges in the graph to generalize this approach to other codon or nucleotide scoring schemes. Also, we assume that LCAI(*u, v*) = LCAI(*v, u*). That is, our weight matrix represents bi-directed edges combining the left-to-right direction and the right-to-left direction. Default weights are calculated using the human codon frequencies in the Kazusa database [27]. Terms used for codon graph access are summarized in Table 3 and depicted visually in Figure 1C.

**Table 3:**
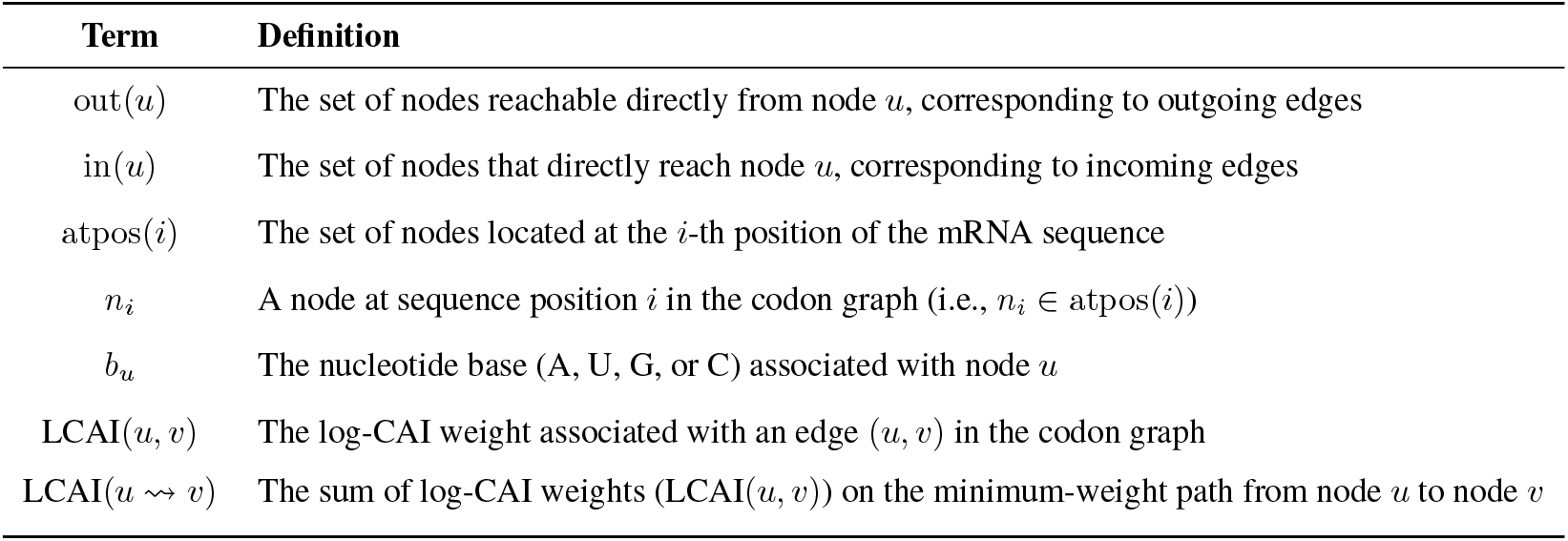
Definitions for Codon Graph Access.

| Term | Definition |
| --- | --- |
| $\text{out}(u)$ | The set of nodes reachable directly from node $u$, corresponding to outgoing edges |
| $\text{in}(u)$ | The set of nodes that directly reach node $u$, corresponding to incoming edges |
| $\text{atpos}(i)$ | The set of nodes located at the $i$-th position of the mRNA sequence |
| $n_i$ | A node at sequence position $i$ in the codon graph (i.e., $n_i \in \text{atpos}(i)$) |
| $b_u$ | The nucleotide base (A, U, G, or C) associated with node $u$ |
| $\text{LCAI}(u, v)$ | The log-CAI weight associated with an edge $(u, v)$ in the codon graph |
| $\text{LCAI}(u \rightsquigarrow v)$ | The sum of log-CAI weights ($\text{LCAI}(u, v)$) on the minimum-weight path from node $u$ to node $v$ |

We define LCAI(*n*_*i*_ ⇝ *n*_*j*_) as the sum of log-CAI weights on the minimum-weight path from node *n*_*i*_ to node *n*_*j*_. For simplicity, we assume *i* ≤ *j* and let LCAI(*n*_*i*_ ⇝ *n*_*j*_) = ∞ otherwise. Let LCAI(*u* ⇝ *v*) = ∞ if there is no path. LCAI(*u* ⇝ *v*) can be computed ahead of time and stored in a table. This can be done using dynamic programming as demonstrated in Equation (4).

## 3 New mRNA Algorithm

This section describes our new algorithms. We start by explaining the details of our dynamic programming method. Our algorithm uses a codon graph, which is similar to existing mRNA folding algorithms. However, we must modify the dynamic programming recurrence to enable suboptimal folding. As a result, our new recursions differ from prior algorithms. Next, we explain how suboptimal folding can be done using our modified dynamic programming recursions. Then, we discuss how our method can incorporate UTRs, then how it be executed using multiple threads, and finally the strategies we employ to control local structures.

### 3.1 New Dynamic Programming Algorithm

Our dynamic programming recursions differ from existing mRNA folding methods in the literature because they are unambiguous (Figure 2). This is important for suboptimal folding. Ambiguous recursions can describe a single possible solution in multiple different ways, and these will lead to duplicates during suboptimal folding. This has been observed in RNA folding algorithms [10], and unambiguous grammars must be used for RNA suboptimal folding [44] and partition function calculation [24]. The recursions for CDSfold are ambiguous [37], as they follow the Zuker-Stiegler recursions. The full recursions for LinearDesign are not formally defined, but the simplified recursions provided are ambiguous [45], and since they do not implement suboptimal folding, we assume their full recursions are also ambiguous.

**Figure 2.**
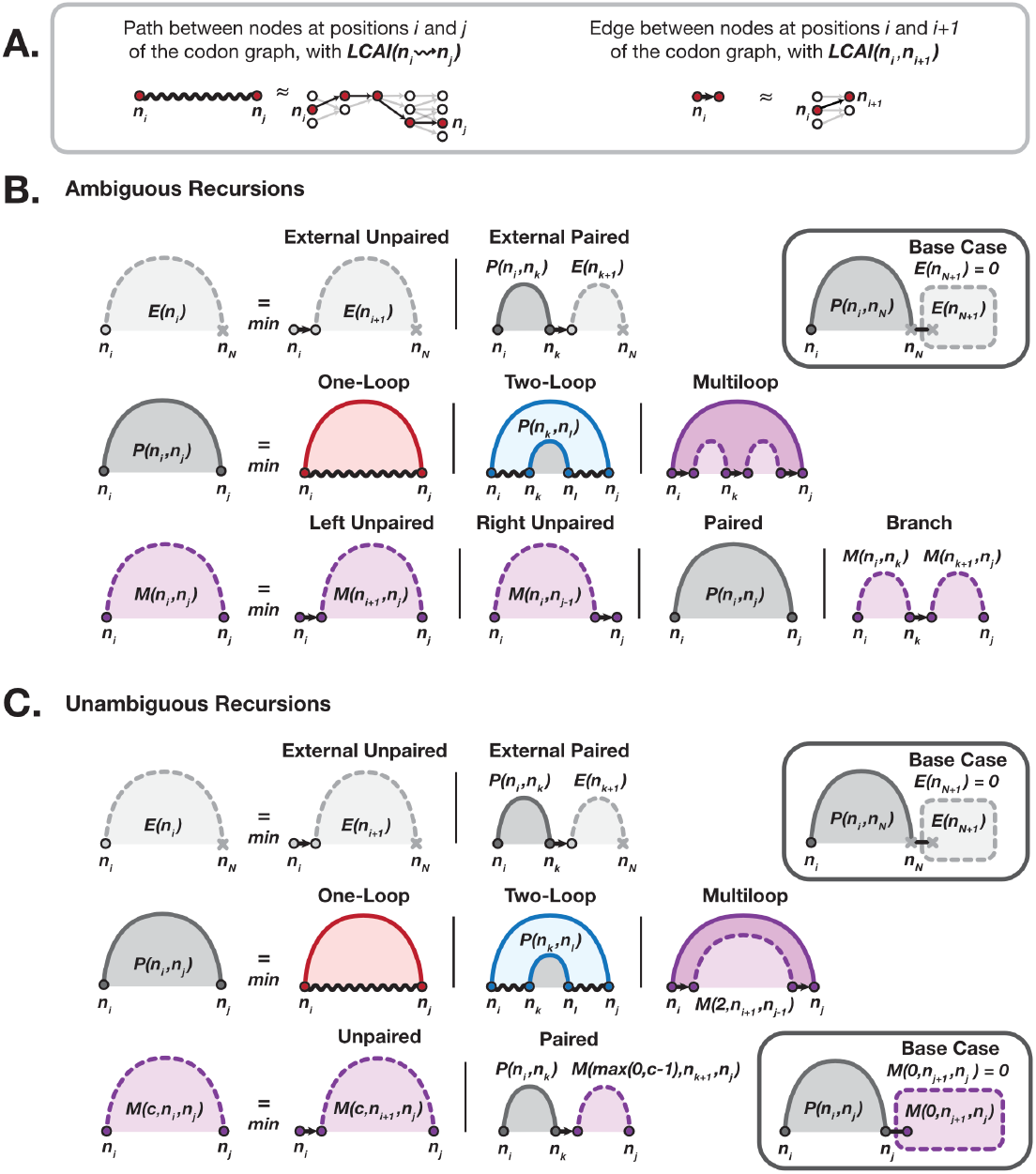
mRNA folding recursions. (A) mRNA folding algorithms extended the RNA folding recursions to work over a codon graph, referencing each nucleotide with a pointer to a node in the graph (e.g. *n*_*i*_ or *n*_*j*_). Paths between positions are replaced with paths along edges in the graph, which incorporate LCAI weights. (B) Existing mRNA folding algorithms follow the standard Zuker-Stiegler recursions. Each case of the recursions is depicted with a Feynman-style diagram [32]. LCAI values are omitted from this visualization for clarity. A solid arc represents a base-pair between nodes at two positions in the sequence. For example, *P* (*n*_*i*_, *n*_*j*_) is represented as a solid arc between node *n*_*i*_ and *n*_*j*_. A dashed arc indicates that nodes at two positions that may or may not be paired. For example, *M* (*n*_*i*_, *n*_*j*_) is represented as a dashed arc between *n*_*i*_ and *n*_*j*_ since nodes at these positions are not necessarily paired with each other. A wavy black line segment at the base of the arc indicates that the intervening positions are unpaired. In all other cases, the structure of the intervening positions is not yet determined. Colors correspond to the corresponding loop type for each case: one-loop (red), two-loop (blue), or multiloop (purple). Cases that do not correspond to a specific loop type are colored grey. The base case for *E*(*n*_*N*+1_) is shown to demonstrate how end-to-end base pairing is considered by the recursions. (C) The mRNA folding recursions can be made unambiguous by replacing the multiloop case (purple). These new recursions are otherwise equivalent. The base cases for *E*(*n*_*N*+1_) and *M* (0, *n*_*j*_ + 1, *n*_*j*_) is shown to demonstrate how end-to-end base pairing is considered by the new recursions.

Consider the multiloop fragment dynamic programming function from [41]. It calculates the minimum scoring multiloop structure fragment over all sequences encoded by paths in the codon graph from node *n*_*i*_ to node *n*_*j*_. To do so, it generalizes the recursions used by CDSfold and LinearDesign to operate on a codon graph:

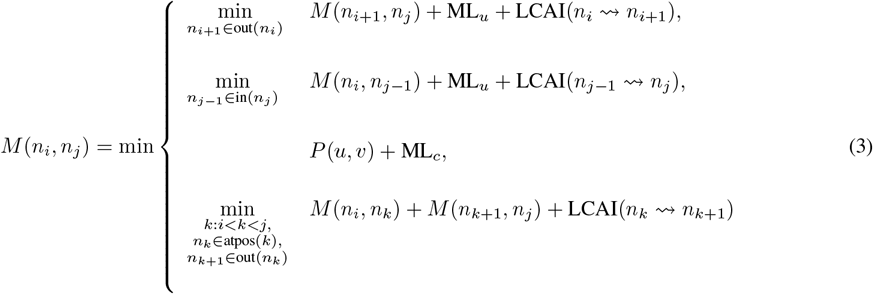

Ambiguous recurrences “double count”. In the case of mRNA folding, this means there at least two distinct ways (and possibly many more) to express the same sequence and structure using the recursions. In Equation (3), we can express a structure that has an unpaired nucleotide on the left (at *i*) and an unpaired nucleotide on the right (at *j*) for a given pair of nodes *n*_*i*_ and *n*_*j*_ in two ways. We could take the first recurrence (the *M* (*n*_*i*+1_, *n*_*j*_) case), then take the second (the *M* (*n*_*i*_, *n*_*j*_ − _1_) case). Alternatively, we could take the second recurrence, then take the first (Figure 3). Both result in positions *i* and *j* being unpaired. This is ambiguous, which means we will see duplicates when enumerating suboptimal results.

**Figure 3.**
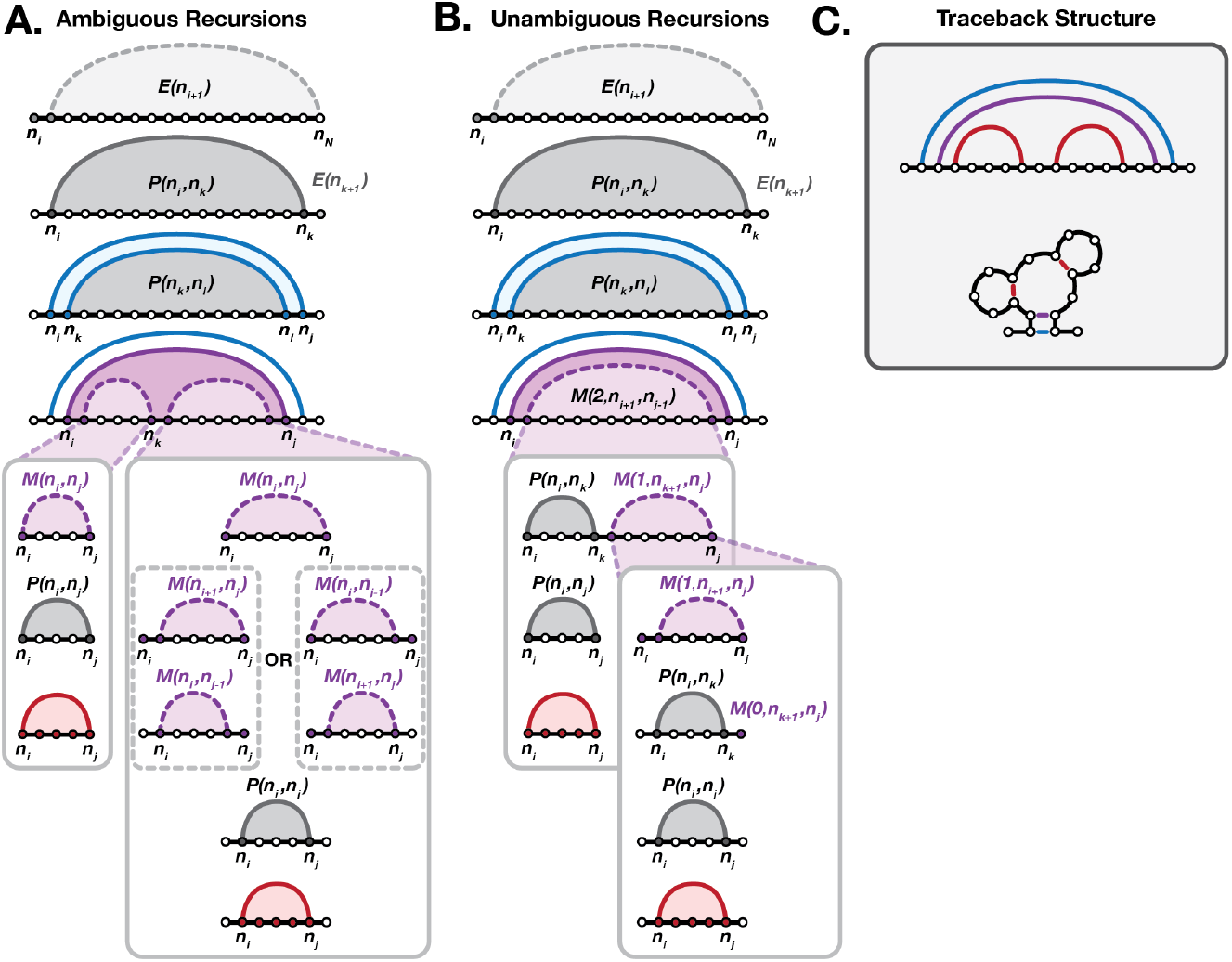
Example multiloop recurrences. (A) The recursion case called at each step of structure prediction is shown for a simple toy example. Filled circles indicate the nodes whose structure is being resolved at the given step. Ambiguous recurrences can generate the same structure via multiple paths through the recurrences. In this example, the two equivalent paths are indicated in dashed grey boxes. (B) Unambiguous recursions can only generate this example structure via a single path through the recurrences. (C) The complete nested secondary structure is the same for both sets of recursions. The structure for this toy example is shown as an arc diagram (left) and its corresponding secondary structure (right), with colors corresponding to the loop type as in Figure 2.

We now describe a new set of recursions that are unambiguous. To begin, LCAI can be precomputed using dynamic programming:

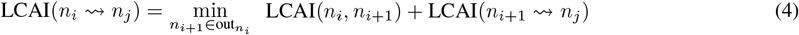

The base case for LCAI(*n*_*i*_ ⇝ *n*_*j*_) occurs when *i* = *j*. In this case, let LCAI(*u* ⇝ *v*) = 0 when *u* = *v* and LCAI(*u* ⇝ *v*) = ∞ when *u*≠ *v*.

The primary change from the ambiguous recursions is to the *M* function. Compare Equation (3) to Equation (6). The last case in the *P* function has been modified to use the new *M* function, and *E* is unchanged.

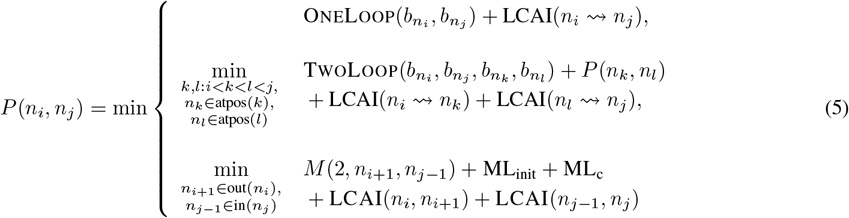

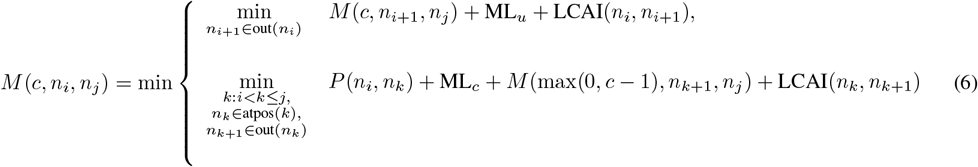

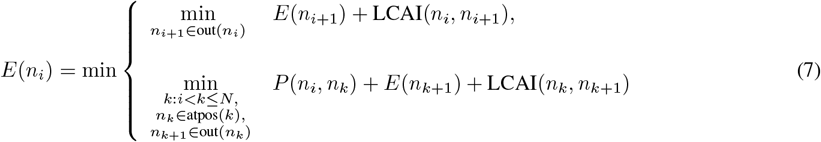

*P* (*n*_*i*_, *n*_*j*_) is the minimum CAIMFE over all sequences with paths starting at node *n*_*i*_ and ending at node *n*_*j*_. Also, it is assumed that sequence locations *i* and *j* are base paired. The only base case is when 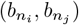 is not a valid bonding pair of bases in the energy model (e.g.,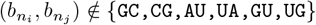).

*M* (*c, n*_*i*_, *n*_*j*_) is the minimum CAIMFE over all sequences with paths starting at node *n*_*i*_ and ending at node *n*_*j*_. Also, only structures that are multiloop fragments are considered, and the multiloop fragment must contain at least *c* closing pairs (i.e., calls to *P* (*u, v*)). The base cases are *M* (0, *n*_*i*_, *n*_*j*_) = 0 for the empty fragment (*i* > *j*), but *M* (*c* > 0, *n*_*i*_, *n*_*j*_) = ∞ for the empty fragment when more than zero closing pairs are required (*i* > *j* ∧ *c* > 0). Since the *i* > *j* base case rules require *i* > *N* to be defined where *N* is the length of the sequence in nucleotides, we construct a special “end node” *ω* in codon graph at index *N* + 1. There are edges from all nodes at position *N* to *ω*. This construction assumes that sequence positions are labeled left to right from 1 to *N*.

The parameter *c* in *M* (*c, n*_*i*_, *n*_*j*_) is used to ensure that valid multiloop fragments containing only a sufficient number of branches are considered. Since a multiloop must have at least 3 branches (otherwise it is a one or two-loop), we can use *c* = 2 when we close a multiloop as the closing pair contributes one branch. To this end, *c* ∈ { 0, 1, 2 } is assumed in our algorithm.

*E*(*n*_*i*_) is the minimum CAIMFE over all sequences with paths starting at node *n*_*i*_ and going to the rightmost end of the codon graph. The base case for *E* is similar to those from *M*. We define *E*(*ω*) = 0 for the empty suffix. The base case is thus *E*(*n*_*i*_) = 0 when *i* = *N* + 1.

The definition and base case for each table in the unambiguous recursions is summarized in Table 4.

**Table 4:**
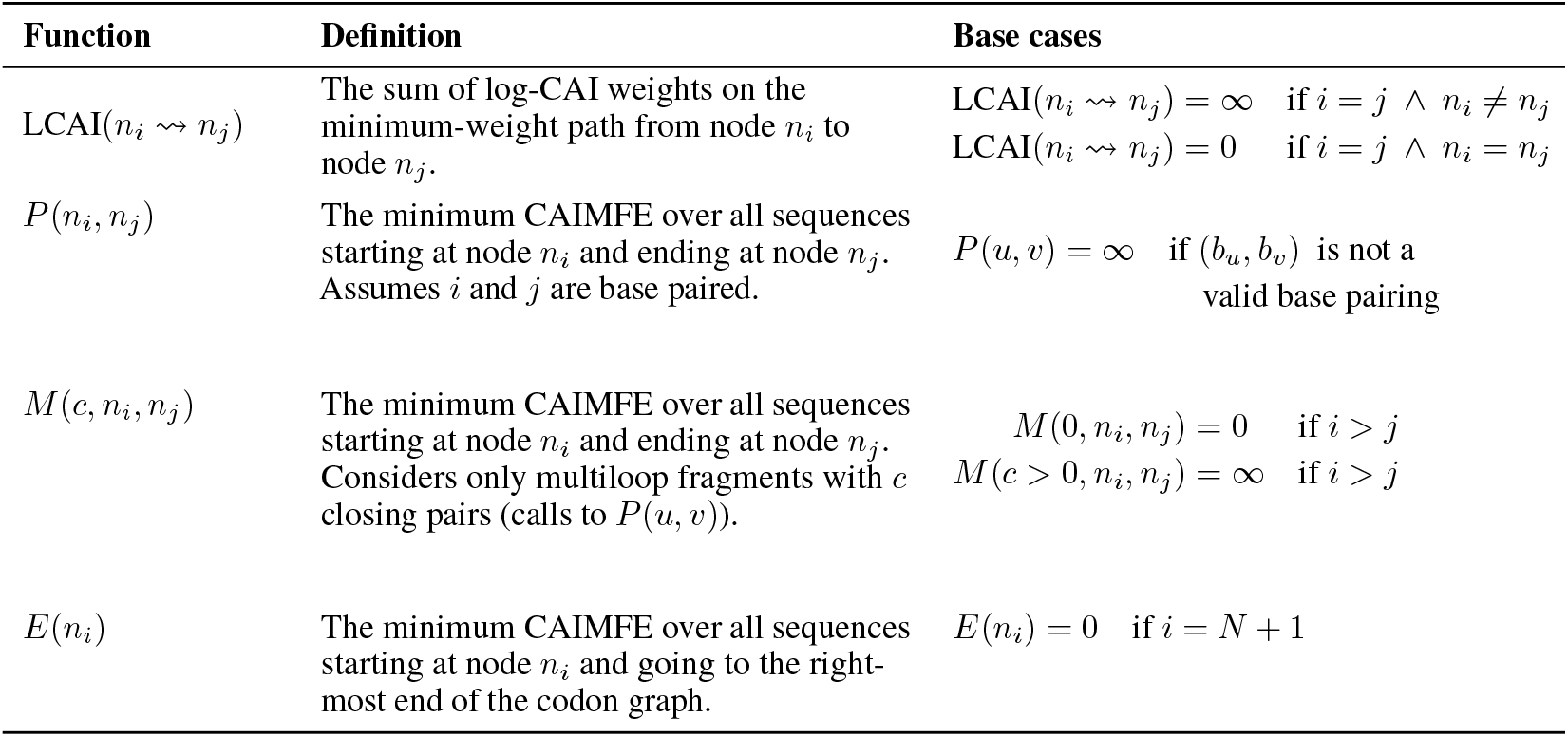
Definitions and base cases for unambiguous dynamic-programming recurrences.

| Function | Definition | Base cases |
| --- | --- | --- |
| $\text{LCAI}(n_i \rightsquigarrow n_j)$ | The sum of log-CAI weights on the minimum-weight path from node $n_i$ to node $n_j$. | $\text{LCAI}(n_i \rightsquigarrow n_j) = \infty$ if $i = j \wedge n_i \neq n_j$<br>$\text{LCAI}(n_i \rightsquigarrow n_j) = 0$ if $i = j \wedge n_i = n_j$ |
| $P(n_i, n_j)$ | The minimum CAIMFE over all sequences starting at node $n_i$ and ending at node $n_j$. Assumes $i$ and $j$ are base paired. | $P(u, v) = \infty$ if $(b_u, b_v)$ is not a valid base pairing |
| $M(c, n_i, n_j)$ | The minimum CAIMFE over all sequences starting at node $n_i$ and ending at node $n_j$. Considers only multiloop fragments with $c$ closing pairs (calls to $P(u, v)$). | $M(0, n_i, n_j) = 0$ if $i > j$<br>$M(c > 0, n_i, n_j) = \infty$ if $i > j$ |
| $E(n_i)$ | The minimum CAIMFE over all sequences starting at node $n_i$ and going to the right-most end of the codon graph. | $E(n_i) = 0$ if $i = N + 1$ |

In general, for *P*, *M*, and also LCAI, we assume the given starting and ending nodes (*n*_*i*_, *n*_*j*_) are in left-to-right order. That is, *i* ≤ *j*. It may be helpful to assume that any states where *j* < *i* have a value of ∞ except where otherwise specified for base cases. However, our algorithm never considers these cases, except as base cases, so they could also be left undefined.

The full-sequence optimal CAIMFE value is obtained by 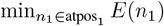.

The time and space complexity of the unambiguous dynamic programming algorithms are *O*(*N*^3^) and *O*(*N*^2^) respectively where *N* denotes the length of the sequence in nucleotides. This is the same as the exist ambiguous versions. The derivation of the complexity is given in the Supplementary Materials.

### 3.2 Suboptimal mRNA Folding

Suboptimal RNA folding is a widely used strategy for analyzing RNA sequences [19, 18, 30, 44, 46]. While there are several methods, for simplicity we define suboptimal RNA folding here as picking a free energy change threshold *t*, and finding all structures whose energy score is between the MFE and MFE + *t*. A useful way of thinking about this is that RNA suboptimal folding generalizes MFE folding to find not just the argmin structure, but a sample of structures near the argmin. For the sake of rigor we provide a formal definition. Let *S*(*π*) be the set of all valid structures for an RNA sequence *π*. An RNA suboptimal folding algorithm finds a set of structures {*s* ∈ *S*(*π*): MFE(*π*) ≤ Δ*G*(*s*|*π*) ≤ MFE(*π*) + *t*}.

Our new suboptimal mRNA folding method generalizes this notion to the mRNA folding problem. To understand how, it is useful to reframe the definition of the mRNA folding problem. Previously, we defined it as finding the sequence with minimum CAIMFE in Equation (2). This definition would imply that suboptimal mRNA folding seeks to find a set of sequences whose MFE values are close to the minimum. However, the dynamic programming recursions implicitly consider all sequences and structures, including non-MFE structures, so it is difficult to only sample sequence along with their MFE structure. Instead, we can equivalently define the mRNA folding problem as finding the minimum sequence-structure pair over all possible sequence-structure pairs for a protein. We defined the set of valid sequences for a protein as CDS(*α*), and we defined the set of valid structures for an RNA sequence as *S*(*π*). Let SeqStr(*α*) = ∪_*π*∈CDS(*α*)_ { ⟨*π, s* ⟩: *s* ∈ *S*(*π*) } be the set of all valid sequence-structure pairs. An equivalent definition to Equation (2) is then:

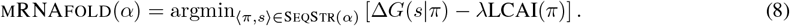

From this we can see that a natural generalization of suboptimal folding is to find a sample of sequence-structure pairs near the minimum. Note that these may not have unique sequences (or structures). That is, two sequence-structure pairs ⟨*π, s*⟩ and ⟨*π*′, *s*′⟩ may have either *π* = *π*′ or *s* = *s*′, but not both.

A further conceptual issue that must be addressed is the choice of threshold *t*. Wuchty’s algorithm, the first complete suboptimal RNA folding algorithm, found all structures within a window of the MFE defined by an energy threshold *t* [44]. This is natural, since free energy change is an interpretable quantity, so finding all structures within 1 *kcal/mol* (for instance) is well defined. For mRNA suboptimal folding, we could seek to find all sequence-structures within a threshold *t* of the CAIMFE (see Equation (1)). The issue is that the combination of energy and CAI used in CAIMFE is no longer just a free energy change and is difficult to interpret. For instance, the meaning of the threshold changes as *λ* changes. Instead, we adopt a modified objective for suboptimal mRNA folding. We take the top-*K* sequence-structure pairs. So, let *L* = EnergyCAISort(SeqStr(*α*)) denote the list of all valid sequence-structure pairs sorted in ascending order by their combined free energy and log-CAI values (Δ*G*(*s* | *π*) − *λ*LCAI(*π*)). We define suboptimal mRNA folding as finding the first *K* entries of *L*.

We are not aware of a similar approach being used for RNA folding or mRNA folding in the literature. However, we note that a top-*K* approach for RNA folding (but not mRNA folding) appears in the `memerna` software package [6], but is unpublished at time of writing.

#### 3.2.1 Priority Queue-based Tracebacks

Solutions are extracted from dynamic programming tables by way of a traceback (also often called backtracking). This process involves traversing the path through the various recursions contributing to an optimal solution to reconstruct the choices involved. In RNA folding, this involves finding the set of base pairs in the structure. In mRNA folding, it involves finding the bases comprising the sequence as well as the pairs comprising the structure.

Since we want to find the top-*K* tracebacks, we use a priority queue of partial tracebacks. A partial traceback 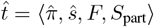 has several parts: a partially complete sequence 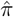 a partially complete structure *ŝ*; a set of states *F* that represent the frontier of the sequence-structure that is not yet determined (Figure 4A); a partial score *S*_part_ for the so far determined sequence-structure. Each state in the frontier *F* corresponds to a location in a dynamic programming table the traceback algorithm should process next (e.g., *P* (3, 4) or *E*(6) as in Figure 4B). These represent the parts of the sequence and structure not yet constructed, and therefore not present in 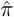 or *ŝ*. The partial score *S*_part_ sums over energies (e.g.,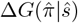) and CAI contributions (e.g.,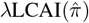) that are already determined. In principle, *S*_part_ could be calculated from 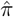 *ŝ*, and *F*. For clarity, we explicitly include *S*_part_ in 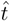.

**Figure 4.**
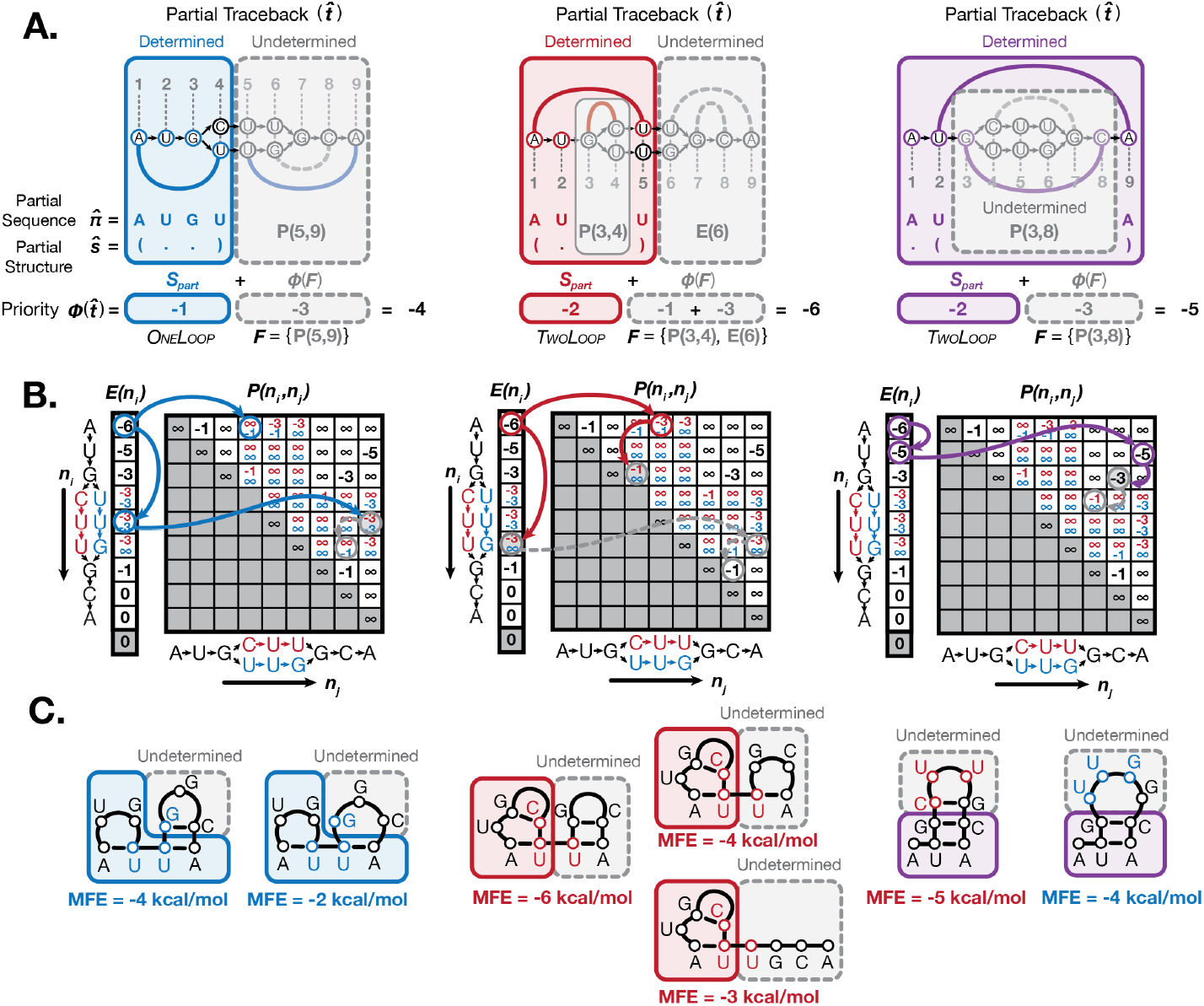
Suboptimal folding partial tracebacks. We use a set of simplified energy terms purely for demonstration purposes: we set the energy of a OneLoop to −1 kcal/mol and the energy of a TwoLoop to −2 kcal/mol and no CAI contributions (e.g., the LCAI table) are used. We also only allow canonical GC and AU base pairs and exclude multiloops. (A) Three example partial tracebacks 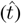 are shown for a simple codon graph. The determined portion of the sequence-structure is indicated in blue, red, or purple, while the undetermined portion(s) of the sequence-structure are in grey. The partially complete sequence 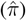 and structure (*ŝ*) are shown below each traceback along with the priority score. (B) The traceback path through the dynamic programming tables is shown for each example. The *F* states are highlighted in grey. Dashed lines indicate the minimum MFE path for the remainder of the traceback. (C) The possible structures corresponding to each partial traceback are shown, along with their final MFE values. Determined portions of the sequence-structure are highlighted in blue, red, or purple, while possible structures for the undetermined portions of the sequence-structure are highlighted in grey. The priority score in (A) and traceback in (B) corresponds to the best scoring sequence-structure (lowest MFE) of these options, though the complete suboptimal traceback could generate any of them.

Each partial traceback has a priority that determines its order in the priority queue. The priority queue is ordered by lowest priority first. Let 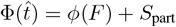 denote the priority function. The function *ϕ*(*F*) = _*f*_ ∈_*F*_ Lookup(*f*) computes the minimum possible combined energy and CAI value for the undetermined parts of the sequence-structure by examining the set of states remaining. The Lookup(*f*) function denotes looking up the value stored in the dynamic programming tables for a table location *f*. The idea is that *S*_part_ is the score of the sequence-structure so far, and *ϕ*(*F*) computes an optimistic “best possible” score for the as yet undetermined parts of the sequence-structure. This is equivalent to the CAIMFE for the most optimal sequence-structure that can still form, given the partially determined sequence-structure (Figure 4C). Three examples of partial tracebacks and their priority score calculation are shown in Figure 4, with simplified energy terms for demonstration purposes.

To generate the top-*K* tracebacks, we only need to keep the top-*K* partial tracebacks. As such, we limit the size of the priority queue to *K*.

The pseudocode in Algorithm 1 provides an overview of the suboptimal folding algorithm. The logic is broken up into Trace*E*, Trace*P*, Trace*M*, and TraceLCAI. These functions recapitulate the dynamic programming recursions. The same basic principles are used in each to translate from dynamic programming to traceback.

It is worth pointing out that the LCAI table is included in the traceback. It is easy to miss the significance of this, which is that it allows us to recover the sequence identity of stretches of unpaired nucleotides.

The suboptimal folding pseudocode makes use of 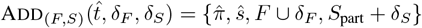, which adds the states subset *δ*_*F*_ to *F* and adds the combined energy and log-CAI score *δ*_*S*_ to *S*_part_. Similarly, the 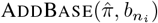 adds the base 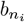 to position *i* in the partial sequence 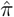, and AddPair(*ŝ, i, j*) inserts a base pair (*i, j*) to a partial structure *ŝ*.

##### Algorithm 1 Suboptimal traceback algorithm

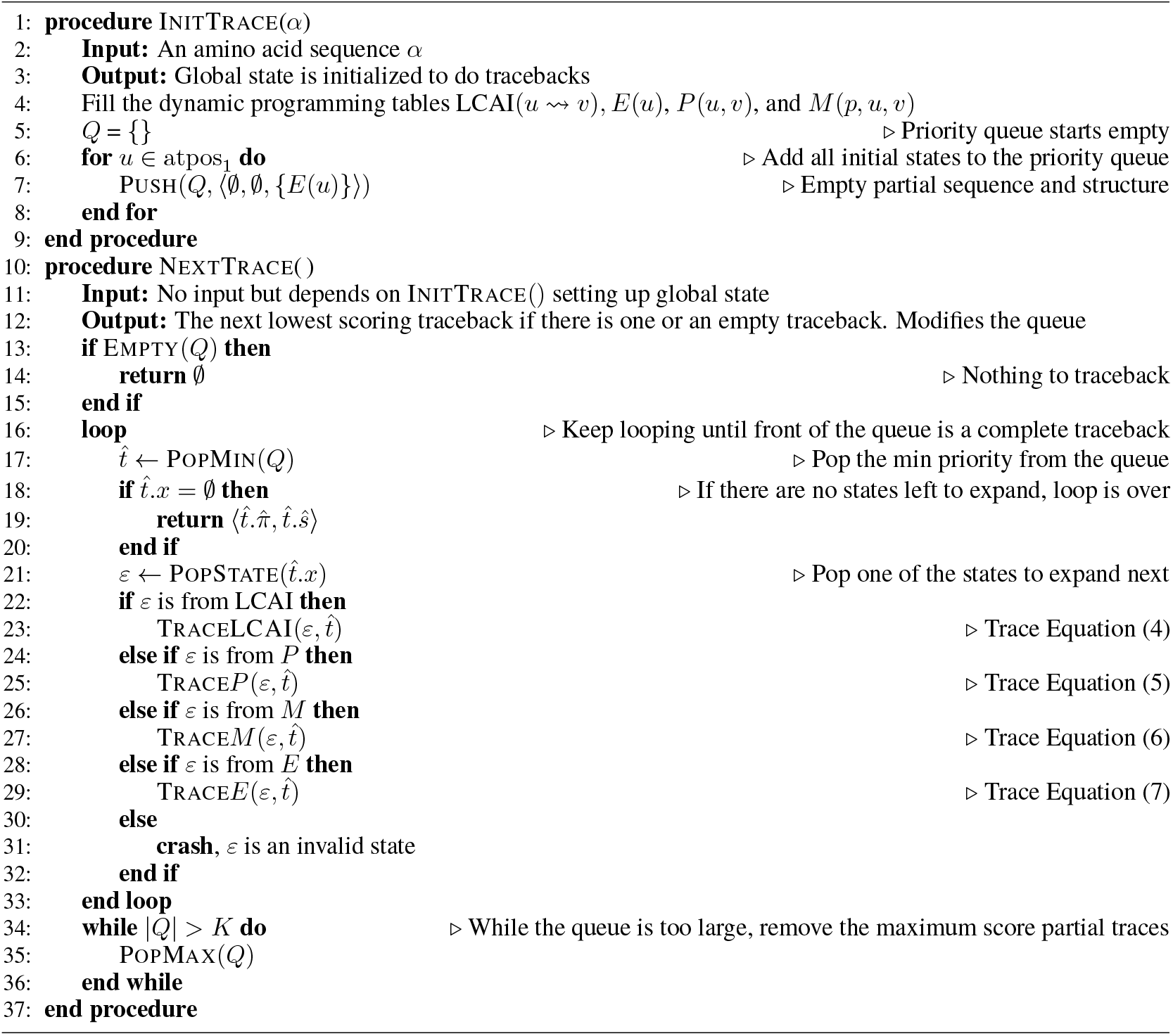

##### Algorithm 2 Traceback for LCAI(*n*_*i*_ ⇝ *n*_*j*_). See Equation (4).

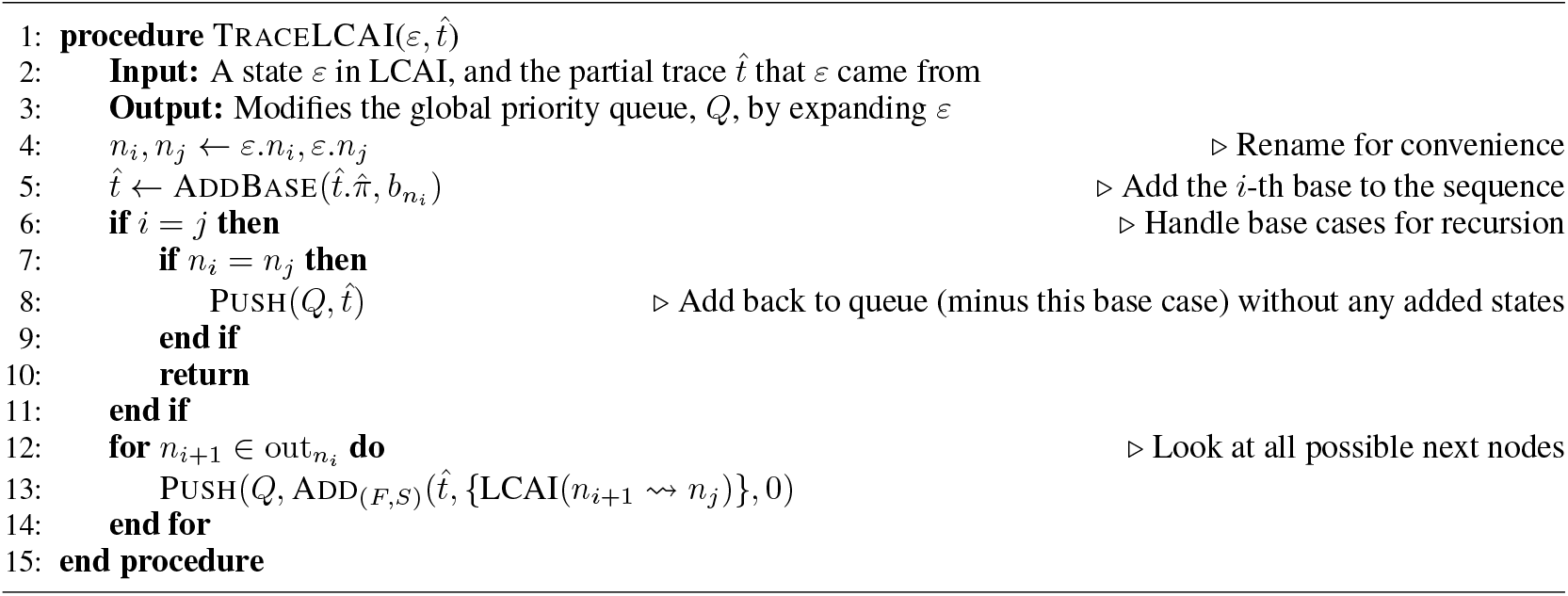

##### Algorithm 3 Traceback for *P* (*n*_*i*_, *n*_*j*_). See Equation (5).

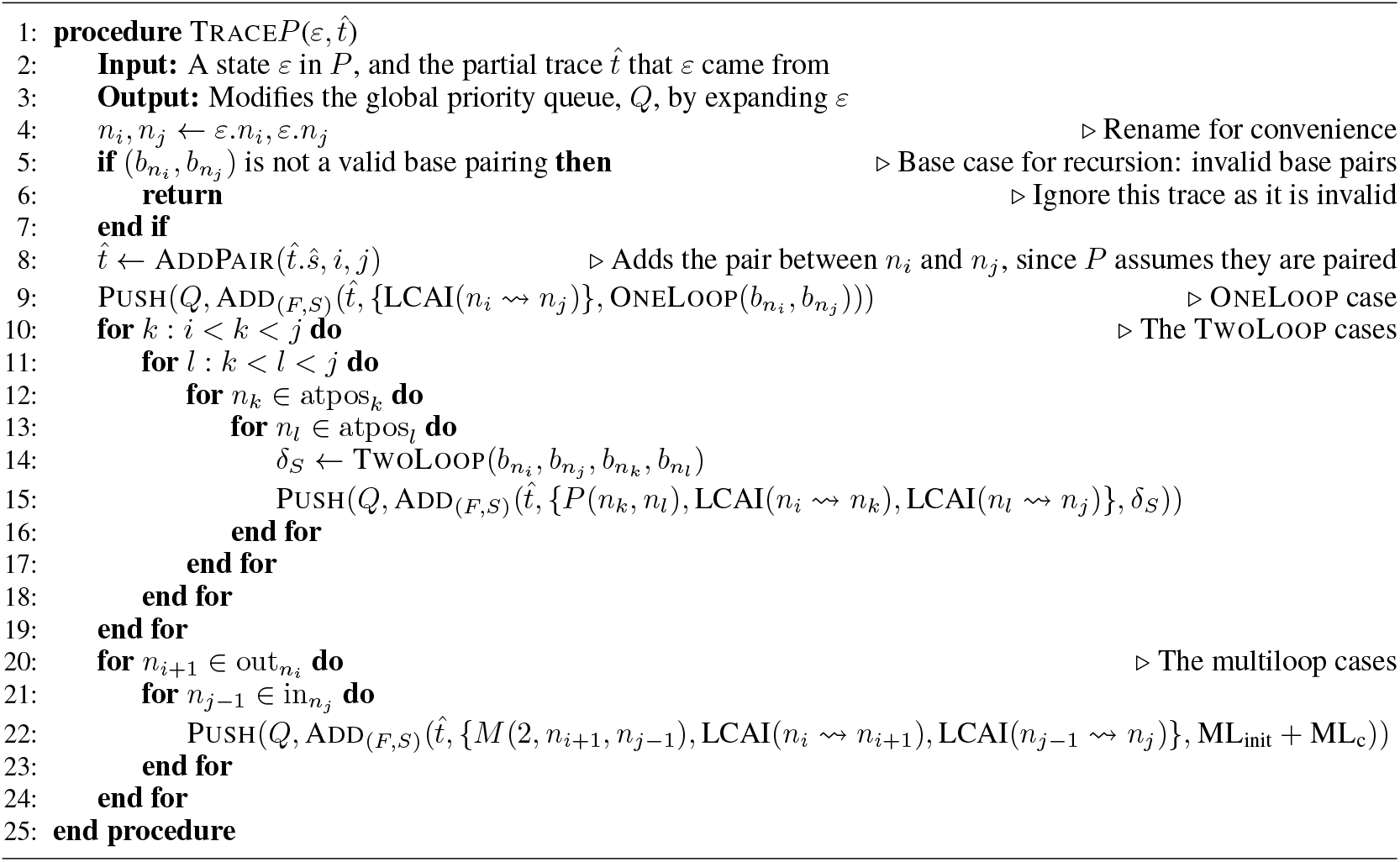

To generate the top-*K* tracebacks, one can call InitTrace(*α*) followed by *K* calls to NextTrace(). The pseudocode assumes that *K* is a global constant. However, we observe that the PopMax(*Q*) loop at the end of NextTrace() could be removed. This allows the algorithm to work as a lazy iterator that can generate any number of tracebacks in order of optimality stopping whenever the user requires.

The pseudocode for Trace*M* and Trace*E* is provided in the Supplementary Materials. It follows the same principles as above is can be derived by applying them to Equation (6) and Equation (7). The complexity analysis for the suboptimal folding algorithm is also given in the Supplementary Materials. The total worst case complexity is thus *O*(*KN*^3^ + *KN*^2^ log *K*) to extract the top-*K* suboptimal tracebacks for a sequence comprising *N* nucleotides. This may seem high, but in practice it is usually much better. This analysis assumes all tracebacks are independent, but often they share many partial traces. Also, often a traceback does not require processing *O*(*N*) partial traces. That is, many traces overlap in sequence and structure.

#### 3.2.2 Increasing Sample Diversity

An issue with suboptimal RNA folding algorithms is that they tend to sample many similar structures. A large sample of structures may contain only a few distinct structural clusters and many small variations on similar structures. This issue is even worse for suboptimal mRNA folding. Not only do we see many small variations on similar structures, but also small variations on similar sequences. We employ a stochastic suboptimal folding method to address this issue. The method modifies the traceback procedure so they have a well defined proportion of randomness.

Recall that the priority of a trace 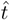 in the priority queue *Q* is calculated using the priority function 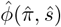. A random variable *r* drawn from a continuous uniform distribution *r* ~ *U* [0, ℛ) can be introduced: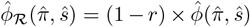. The parameter ℛadjusts the amount of randomness. In this new formula, the priority of any 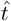 is partly random, and the randomness is proportional to ℛ. For instance, if we set ℛ= 0.5, then the priority is half random, and if we set it to ℛ= 1.0, then it is entirely random. In essence, we use random proportional down-scaling with adjustable proportionality.

### 3.3 Parallel Folding

Since mRNA folding is computationally intensive, we implemented a parallel version of the dynamic programming algorithm. This requires solving the recursions using a bottom-up table fill. The key insight is that *P* (*n*_*i*_, *n*_*j*_) (see Equation (5)) only depends on *P* (*n*_*k*_, *n*_*l*_) and *M* (*c, n*_*k*_, *n*_*l*_) where *i* < *k* < *l* < *j*. This means that *P* (*n*_*i*_, *n*_*j*_) only needs to access positions in the *P* and *M* tables where *k* > *i* (Figure 5B). Similarly, *M* (*c, n*_*i*_, *n*_*j*_) only depends on calls to the *P* and *M* tables where the indices are ≥ *i* (Figure 5C), and *E*(*n*_*i*_) only depends on positions ≥ *i* in the *P* and *E* tables (Figure 5D). Finally, LCAI(*n*_*i*_ ⇝ *n*_*j*_) only references positions in LCAI with indices > *i* (Figure 5A).

**Figure 5.**
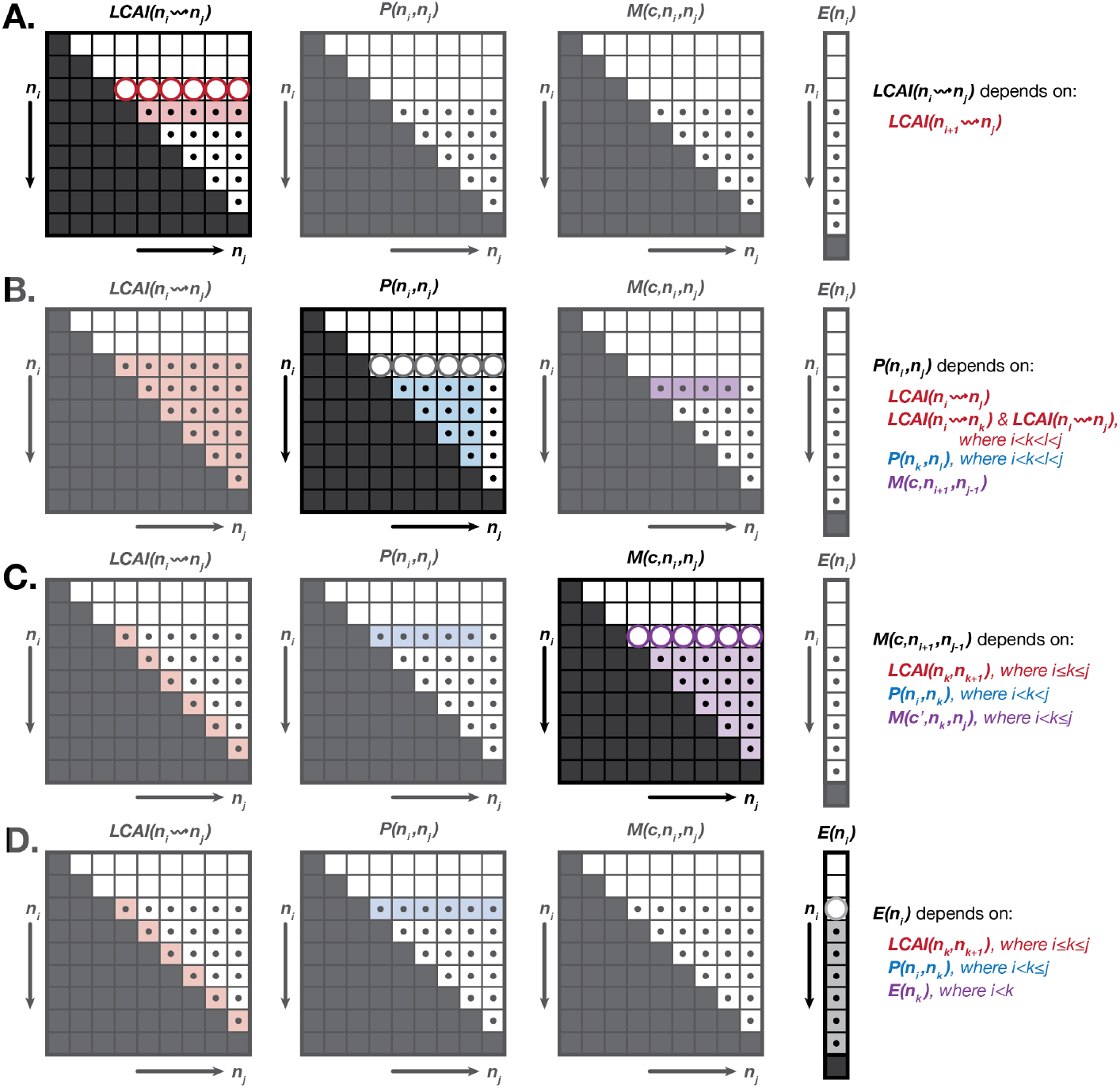
Bottom-up dynamic programming algorithm. (A-D) The dependencies for a given *n*_*i*_ row in each of the dynamic programming tables are shown. The positions that can be calculated in parallel for the current iteration are indicated by large circles, while black dots indicate positions that have already been populated in previous iterations of the *i* loop. Positions that are required for calculation of the current *n*_*i*_ row are highlighted in the appropriate tables. The tables are filled bottom-up, from *i* = *N* to *i* = 1, and are filled in the order shown (A-D) for each *i* value.

This implies we can fill the tables by iterating backwards through values for *i*. Let us define some functions to use. We use Fill*P* (*i, j*) to denote an algorithm that calculates and stores the solutions for all *P* (*n*_*i*_, *n*_*j*_). We define Fill*M* (*c, i, j*), FillLCAI(*i*), and Fill*E*(*i*) analogously. Our bottom-up algorithm is given in Algorithm 4.

#### Algorithm 4 Outline of the bottom-up table fill routine for the our dynamic programming algorithm.

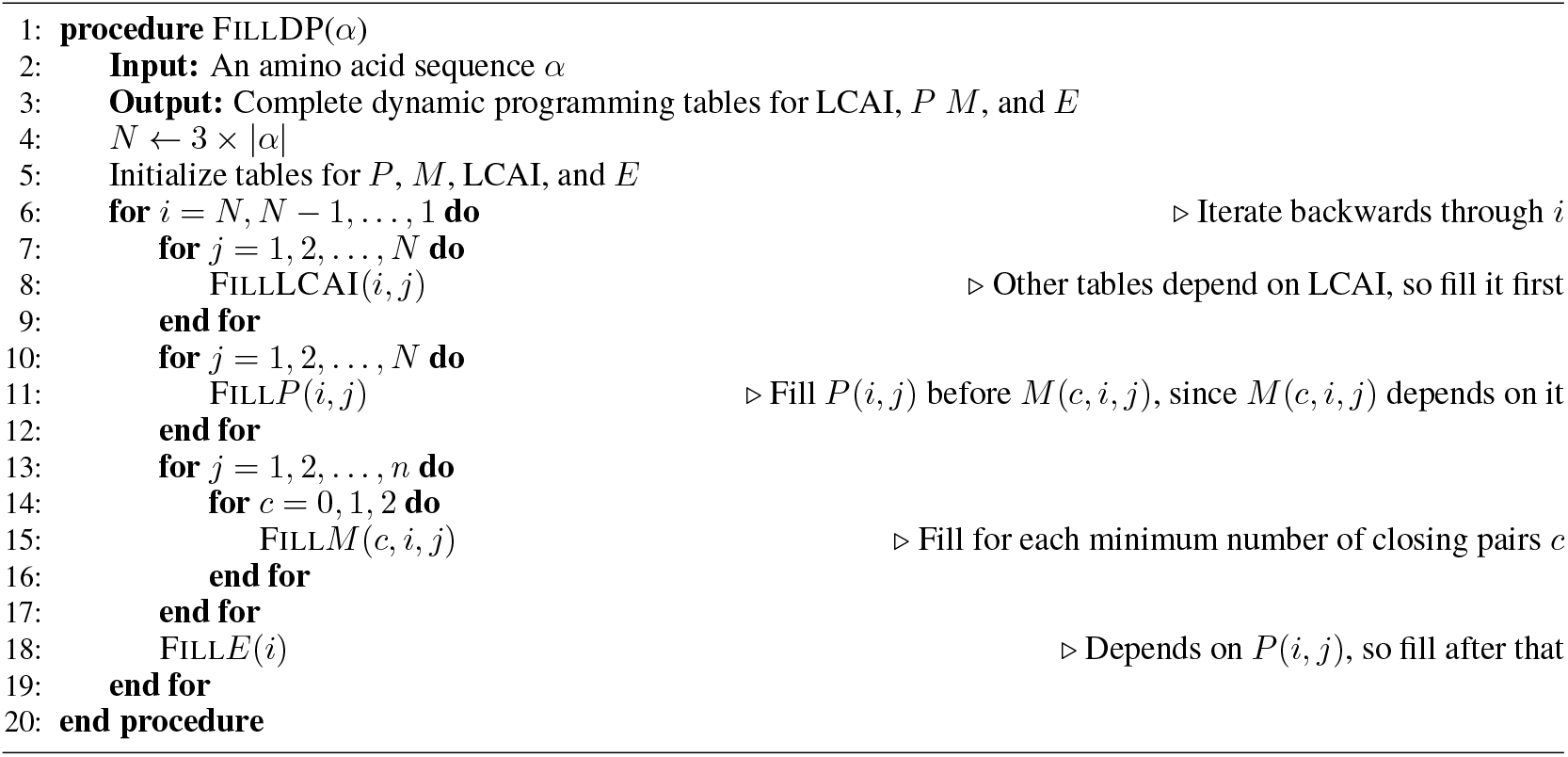

Observe that there are no dependencies between *j* values in Algorithm 4. Specifically, the order of the *j* values in the for loops on lines 8, 11, and 15 are arbitrary. The values for *j* could be executed in any order and it would not affect the correctness of the program. This is not true for the *i* for loop. The upshot of this is that we can execute each *j* for loop in parallel. This reduces the complexity of the algorithm from *O*(*N*^3^) to *O*(*N*^2^), assuming enough processors are available.

### 3.4 Incorporating UTRs

Our algorithm is the first mRNA folding algorithm to holistically incorporate the UTRs into the optimization. This is important since the structure of a sequence may change with the UTRs present. Base pairs may form between the bases in the UTR and the bases in the CDS.

The UTRs are perhaps surprisingly straightforward to incorporate using a codon graph model. We prepend a path representing the nucleotides in the 5’UTR and append a path representing the nucleotides in the 3’UTR (Figure 6).

**Figure 6.**
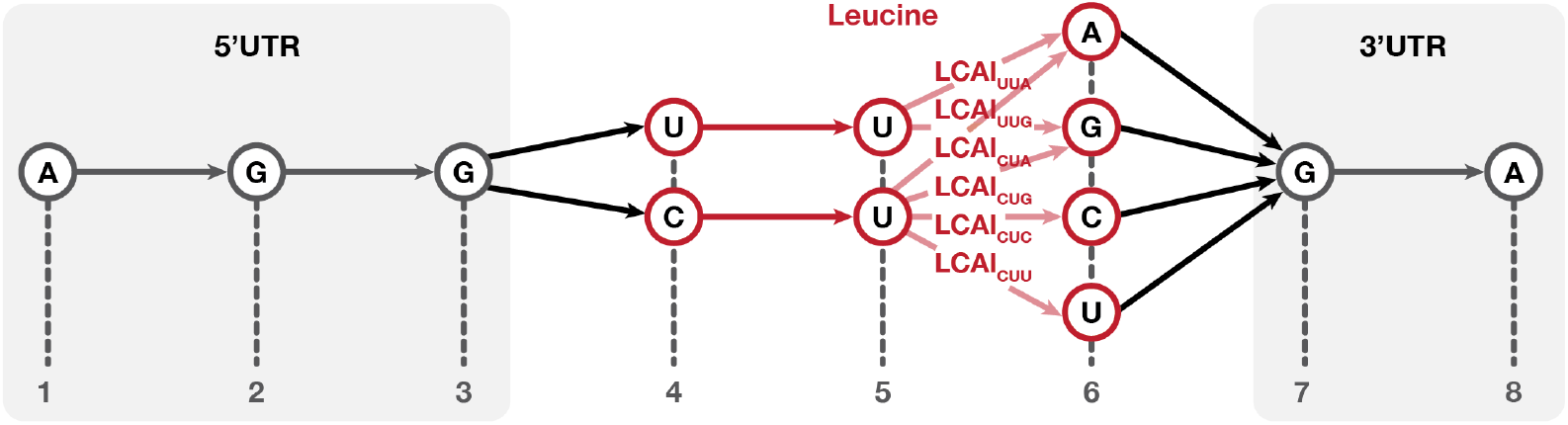
Codon Graph with UTRs. An example codon graph with the 5’UTR “AGG” and the 3’UTR “GA” attached to a leucine codon subgraph. This is demonstrative only, a realistic codon graph would have much longer UTRs, and a much larger CDS component including a start and stop codon.

Recall that, in the codon graph, each path from left-to-right corresponds to a valid sequence. Adding the UTRs as paths ensures that they are a prefix or suffix of each possible path. Note that all the edges in the UTR paths are assumed to have an LCAI weight of zero as the UTRs do not contribute to CAI calculation.

### 3.5 Target Unstructured Regions

It can be desirable to make some regions in the sequence less structured [31]. We extend an approach previously used by CDSfold [37]. This approach’s main weakness is that it is a heuristic that can fail to find a suitable sequence. We use suboptimal folding with diversity to overcome this, as we can sample sequences until a suitable one is found.

Formally, we allow a user to specify a range *R* = [*R*_*i*_, *R*_*j*_] of nucleotides where base pairing should be minimized. This is in contrast with the rest of the sequence, where we generally want to minimize the MFE and therefore increase base pairing. As in the CDSfold heuristic, we introduce a penalty *γ* and set *P*_*γ*_(*n*_*i*_, *n*_*j*_) = *γ* × *P* (*n*_*i*_, *n*_*j*_) where *P*_*γ*_(*n*_*i*_, *n*_*j*_) is the adjusted value we store during dynamic programming if *i* ∈ *R* or *j* ∈ *R*. Typically, *γ* = ∞, which is the case for our experiments. Note that in principle this method can be used to discourage (or encourage) any set of base pairs.

This does not guarantee that the optimized sequence forms no base pairs in *R*. It only ensures that they will not be considered when calculating the MFE. In other words, we ignore the possibility of base-pairs in *R* during mRNA folding (a constrained MFE structure), but the final structure predicted for the same sequence when these base-pairs are allowed may differ (the full MFE structure). The full MFE structure for the optimized sequence may therefore include base-pairs in *R*, but they are less likely. We can modify Equation (8) to reflect this. Define SeqStr*R*(*α, R*) = SeqStr(*α*) \ { sequence-structures with a base pair in *R* }. That is, SeqStr*R*(*α, R*) represents the subset of all possible sequence-structure pairs SeqStr(*α*) that can form without any base-pairs in *R* (constrained MFE structures).

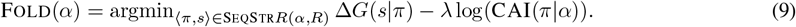

Notice that this does not guarantee that the argmin sequence *π* does not have a full MFE structure with pairs in *R*. It only guarantees that among all sequences, *π* has the most stable constrained MFE structure that avoids pairs in *R*.

To minimize base-pairing, CDSfold used a second optimization stage in which the bases in *R* were modified to reduce structure. We employ a new strategy. We use our new suboptimal folding method to sample a diverse set of sequences and rank them by the amount of structure in *R* without any constraints.

The amount of structure is defined similarly to the measure average unpaired probability [42]. The base-pairing probability table can be used to compute the probability *U*_*i*_ that a nucleotide *i* is unpaired [24, 19]. This requires calculating the base-pairing probability table, which we do using ViennaRNA [18] for each sequence generated by our algorithm. We then compute the average unpaired probability of *R* by 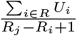 and rank traceback sequences by this measure.

### 3.6 Software Package

The above algorithms are implemented in a new software package called `mRNAfold`. It is available on GitHub at https://github.com/maxhwardg/mRNAfold.

## 4 Experimental Method

### 4.1 Software Benchmarks

We ran benchmarks comparing the runtime and memory usage of `mRNAfold` against established mRNA folding software packages including LinearDesign [45], CDSfold [37], and DERNA [12]. The same methodology from our review of these existing algorithms was followed [41] the key points of which we recapitulate here. Amino acid sequences with lengths between 50aa and 1500aa were used. Two benchmarks were carried out, one using random amino acid sequences, and the other using poly-leucine sequences (i.e., MLLL…). Our `mRNAfold` software package was run both with serial and parallel execution for comparison. All experiments were done on Ubuntu 24.04 LTS and an AMD 7950X 16-core CPU. The GCC 13.3.0 C++ compiler was used to compile all software packages. The most recent GitHub commits at time of writing were used for all software packages: `6a84582`, `f0126ca`, `06f3ee8`, and `fe785ae` for `mRNAfold`, LinearDesign, CDSfold, and DERNA respectively. Our benchmarking code and data are available are https://github.com/maxhwardg/mrna_folding_comparison.

### 4.2 Sequence Design

To demonstrate the application of our new algorithm, we generated optimized eGFP sequences to test two separate biological hypotheses. The first was a highly structured eGFP sequence, which we compared to a high GC content sequence, to test the effect of increased structure on mRNA in-vial stability. The second was a set of eGFP sequences that targeted varying codon optimality and local structure levels to test the impact of these features on in-cell expression.

Selected mRNA sequences were synthesized by *in vitro* transcription with complete substitution of uridine (U) by N1-Methylpseudouridine (m1Ψ) and were co-transcriptionally capped. Each transcript included a 100-nucleotide poly(A) tail encoded in the plasmid template used for transcription.

#### 4.2.1 Design of Sequences for Stability Testing

We designed two eGFP sequences for in-solution stability testing: a high-GC variant and a highly structured variant designed with our new `mRNAfold` algorithm.

To generate the GC-rich eGFP sequence, we selected the synonymous codon with the highest GC content at each position. In cases of ties, codons were chosen based on the highest CAI, calculated using the Kazusa codon usage table [27].

For the MFE-optimized sequence, we used `mRNAfold` with *λ* = 0 (`lambda 0` in the software) to prioritize structural stability over codon preference. We enabled suboptimal folding with randomness ℛ= 0.05 to generate 100 traces (`subopt_randomness 0.05` and `num_subopt_traces 100`). A single candidate was selected from the resulting suboptimal sequence pool for experimental testing. `mRNAfold` uses the energy parameter of ViennaRNA [18], which assume canonical uridine; therefore MFE values were recalculated using Moderna’s proprietary m1Ψ energy parameters.

#### 4.2.2 Design of Sequences for Translation Impact Testing

We generated 18 eGFP sequences with `mRNAfold` to test the impact of local structure and codon optimality on mRNA expression. Broadly, the generated sequences had relatively “low” or “high” levels of codon optimality, along with “low”, “medium”, or ‘high” amounts of structure near the start codon. We refer to each combination of codon optimality and local structure as an “design group” (Figure 9A).

Codon optimality was determined using CAI, calculated using the human codon frequencies from the Kazusa database [27]. For the low CAI sequences, *λ* = 0 (`lambda 0`) was used to reduce the importance of CAI, while for the high CAI sequences *λ* = 10 (`lambda 10`) was used to increase the importance of CAI.

We targeted variable structure in a 20-nucleotide region around the start codon and low structure in the 5’UTR upstream of this region. We define the start codon region as indices 46-65 (one-indexed, inclusive index range), with `AUG` appearing at indexes 58-60.

To vary structure in this region, we used `mRNAfold` to generate sequences using two sets of parameters. For the first run, we targeted low structure in the 5’UTR and start codon region by encouraging unpaired nucleotides in the region *R* = [1, 65] (which corresponds to `encourage_unpaired 0,65`, since the algorithm accepts a zero-indexed, inclusive-exclusive index range). This means we let `mRNAfold` maximize structure in the remainder of the CDS and 3’UTR for the first run. For the second run, we targeted low structure only in the upstream 5’UTR region only by setting *R* = [1, 45] (`encourage_unpaired 0,45`). This means we let `mRNAfold` maximize structure in the start codon region as well as the remainder of the sequence for the second run.

For each set of CAI and structure parameters, we used suboptimal folding with randomness ℛ= 0.1 to generate a diverse pool of 1000 suboptimal traces (`subopt_randomness 0.1`, `num_subopt_traces 100`, `num_subopt_rounds 10`). This generated a diverse set of sequence-structure pairs for each CAI weight.

We then selected sequence-structure pairs with varying amounts of local structure around the start codon. We used Average Unpaired Probability (AUP) as a measure of the extent of global and local structure [42]. Within each CAI group, we selected sequence-structure pairs from the total pool that maintained CAI and global AUP values close to the mean of the pool, while varying AUP in the start codon region *R* = [46, 65]. AUP values were calculated using m1Ψ energy parameters.

The selected sequences for each CAI group include three with AUP values near the minimum for the start codon region (high structure), three with values near the maximum (low structure), and three with values in the middle (medium structure) (Figure 9B). In previous experiments, many mRNAs with very low codon optimality (CAI ≈ 0.6) also exhibited reduced elongation rates, which may have secondarily affected initiation rates [2]. Therefore, we targeted a mid-range codon optimality (CAI ≈ 0.8) for our “low CAI” group and excluded truly low codon optimality constructs from our design.

Details of the procedures used in the experimental assays can be found in the Supplementary Materials.

### 4.3 Statistical Tests

A permutation test was used to determine the statistical significance of the absolute differences between the mean expression level of groups. The area under the curve (AUC) of fluorescence for each construct normalized by the AUC of the standard GFP was used to measure relative expression. The null distribution was estimated by concatenating the data lists from both groups then randomly reassigning group labels (i.e., by permutation). The absolute difference in mean AUC of relabeled groups was calculated for each of 10^6^ samples. The *p*-value was obtained by taking the proportion of values in the null distribution that were greater than or equal to the observed value.

## 5 Results

### 5.1 Software Benchmarks

Figure 7 depicts our benchmarking results. For random sequences (panels A and C), serial `mRNAfold` is faster than all software packages but LinearDesign. Interestingly, `mRNAfold` is initially faster, but they cross at around 500aa. When using all cores, the parallel version of `mRNAfold` is significantly faster than all other algorithms. `mRNAfold` has lower memory usage that all other software packages but CDSfold.

**Figure 7.**
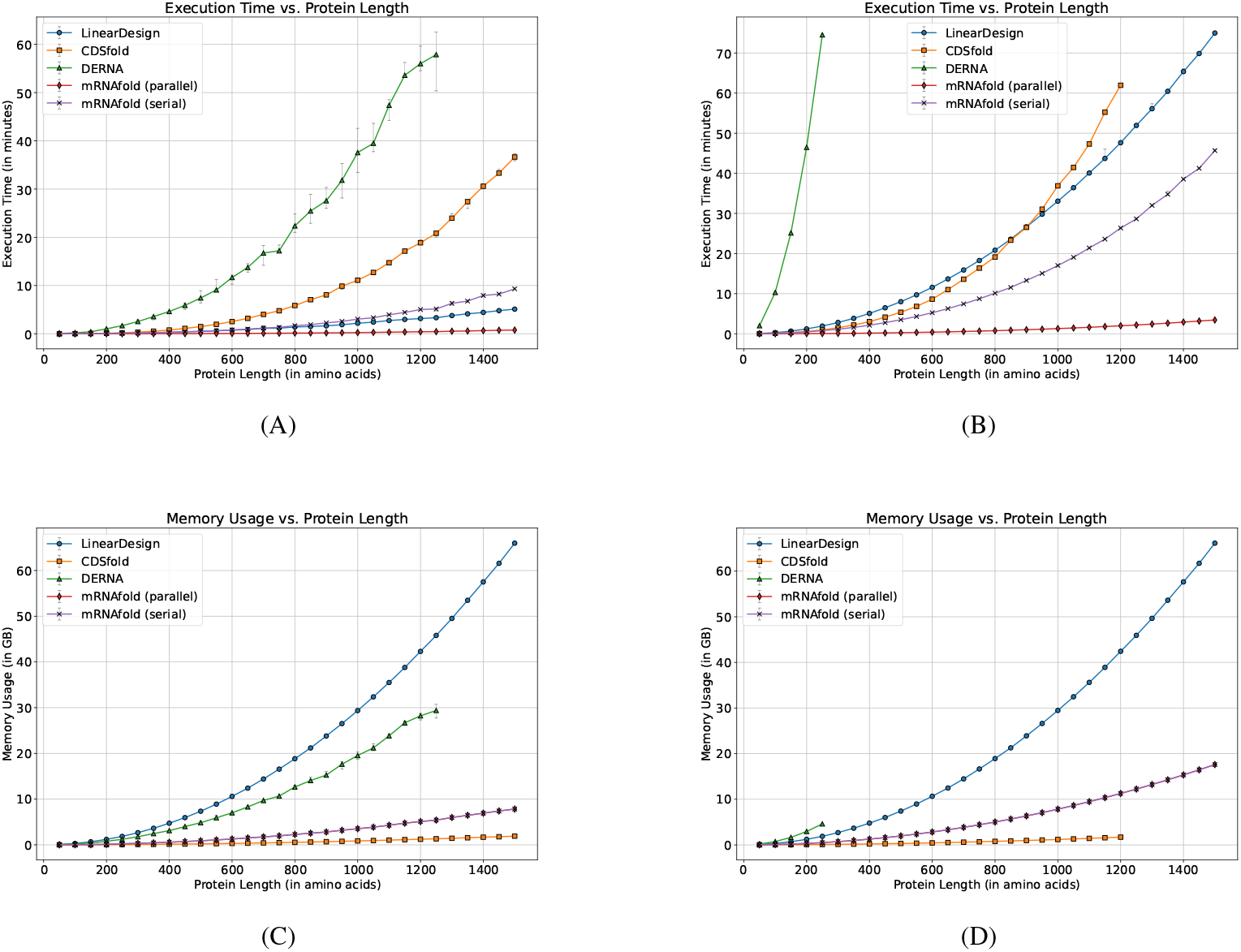
Benchmarking results for time and memory displaying `mRNAfold` and existing mRNA folding software packages. The top row contains execution times for random sequences (A) and for poly-leucine sequences (B). The bottom row contains memory usages for random sequences (C) and for poly-leucine sequences (D). Three runs were done for each data point; the median is displayed with error bars showing the other two results.

For poly-leucine sequences (panels B and D), `mRNAfold` is the faster than all other packages both in serial and parallel modes. The memory usage of all packages is similar to that for random sequences except DERNA, which has significantly increased time and memory requirements.

This comparison comes with some caveats. LinearDesign uses a beam search heuristic that trades off correctness for speed [45], so its speed comes at the cost of sometimes producing suboptimal results. Despite this, `mRNAfold` in serial mode is competitive, being slightly slower for random sequences but slightly faster for poly-leucine sequences. Also, LinearDesign and CDSfold do not use unambiguous recursions, which `mRNAfold` must to enable suboptimal folding. Unambiguous recursions come with a performance cost, since they make some optimizations impossible. Similarly, CDSfold does not incorporate CAI and only minimizes MFE, which means it has more opportunities for optimization. Finally, `mRNAfold`’s parallel mode is capable of using all CPU cores, which other software cannot, so a significant speedup is both expected and observed.

### 5.2 Design for High In-Solution Stability

Increasing mRNA structure has been consistently shown to enhance in-solution stability by reducing susceptibility to hydrolytic degradation [15]. Structural enhancement can be achieved either by increasing GC content or by directly minimizing the minimum free energy (MFE) of folding. We designed two eGFP mRNA sequences: one optimized for high GC content (63% GC, MFE = –439.9 kcal/mol) and another optimized for lower MFE (56% GC, MFE = –616.5 kcal/mol), and evaluated their stability under accelerated degradation conditions at 40^°^C and 25^°^C.

Consistent with previous reports using LinearDesign and RiboTree [45, 15], the MFE-optimized sequence exhibited significantly greater resistance to degradation. At 25^°^C, its half-life increased from an extrapolated 255.9 days (GC-rich sequence) to an extrapolated 318.7 days—a 1.25-fold improvement (Figure 8A). A similar effect was observed at 40^°^C, where the half-life improved from 26.1 to 38 days corresponding to a 1.46-fold enhancement in stability (Figure 8B).

**Figure 8.**
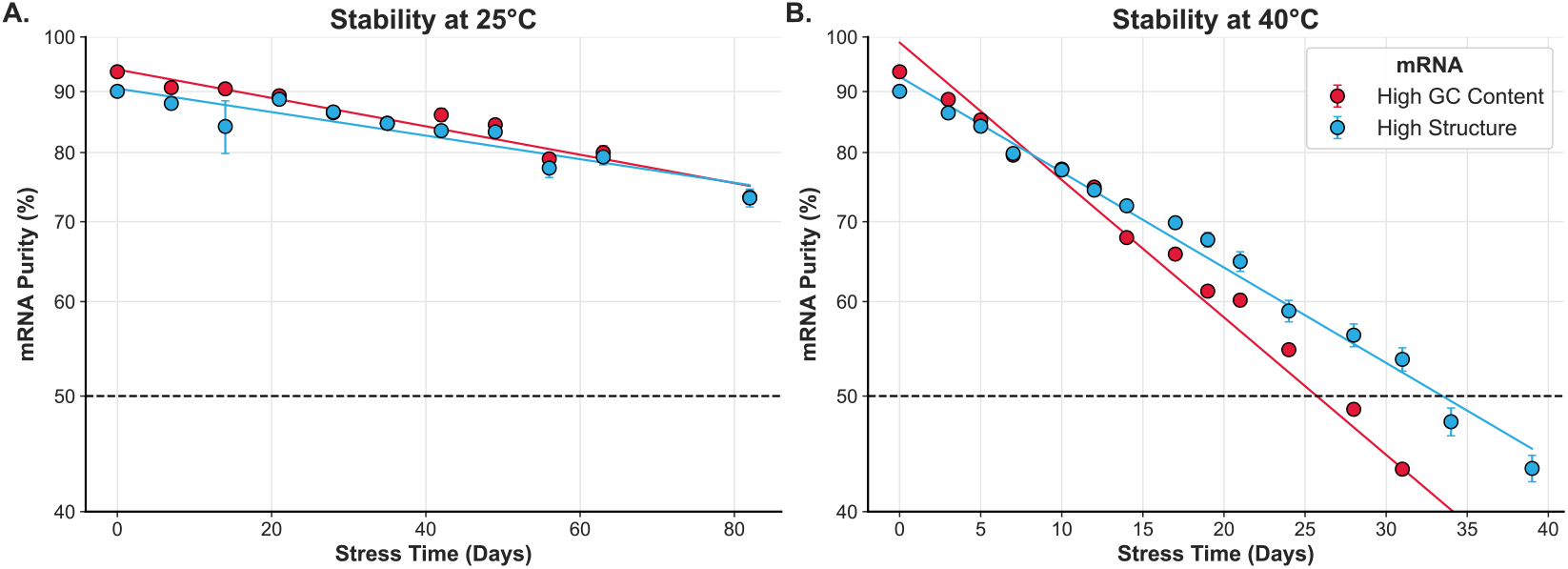
Stability design. (A) In-solution stability of a GC-rich and a high structure mRNA at 25^°^C is shown. (B) The same mRNAs at 40^°^C are shown.

#### 5.2.1 Design for Local Structure Optimization

Our previous work with a set of Luciferase-encoding CDS variants showed a negative correlation between strength of base-pairing in the 5’ region of the RNA (5’UTR plus first 30 nucleotides of the CDS) and protein production, suggesting that low structure in that region can improve initiation rate and thus protein expression [22]. Similarly, in a follow-up study with GFP-encoding mRNAs, high structure in the window surrounding the start codon correlated with low translation initiation rate measurements. However, in that case low initiation rates were associated with high protein production due to higher mRNA half-life [2]. These observations highlight the need for a sequence design algorithm that can modulate RNA structure within specific localized regions while maintaining structure and codon optimality in the rest of the RNA. Here, we used `mRNAfold` to design GFP-encoding mRNA sequences with differing base-pairing probabilities in a 20-nucleotide window surrounding the start codon and Kozak Sequence, while maintaining low predicted structure in the remainder of the 5’UTR and high predicted structure in the CDS. We designed sequence sets with either low or high CAI and measured total protein production for each sequence. We normalized protein production measurements from each mRNA to a control GFP sequence, which we previously designed to have high structure and high CAI throughout the CDS [22, 2].

Figure 9A illustrates the sequence architecture, in which each eGFP encoding mRNA maintained a fixed 5’UTR sequence and a largely conserved downstream CDS (positions 66–909), while the “start codon region” (nt 45–65, flanking the start codon) was engineered to exhibit distinct levels of RNA structure; either low, medium, or high. For each level of start codon region structure, the sequence Hamming distance between the three sequences varied from 0 to a maximum of 167. Although changes to the rest of the CDS were minimized, we could not guarantee identical sequences due to necessary codon substitutions required to modulate structure in the start codon region (Figure 9).

**Figure 9.**
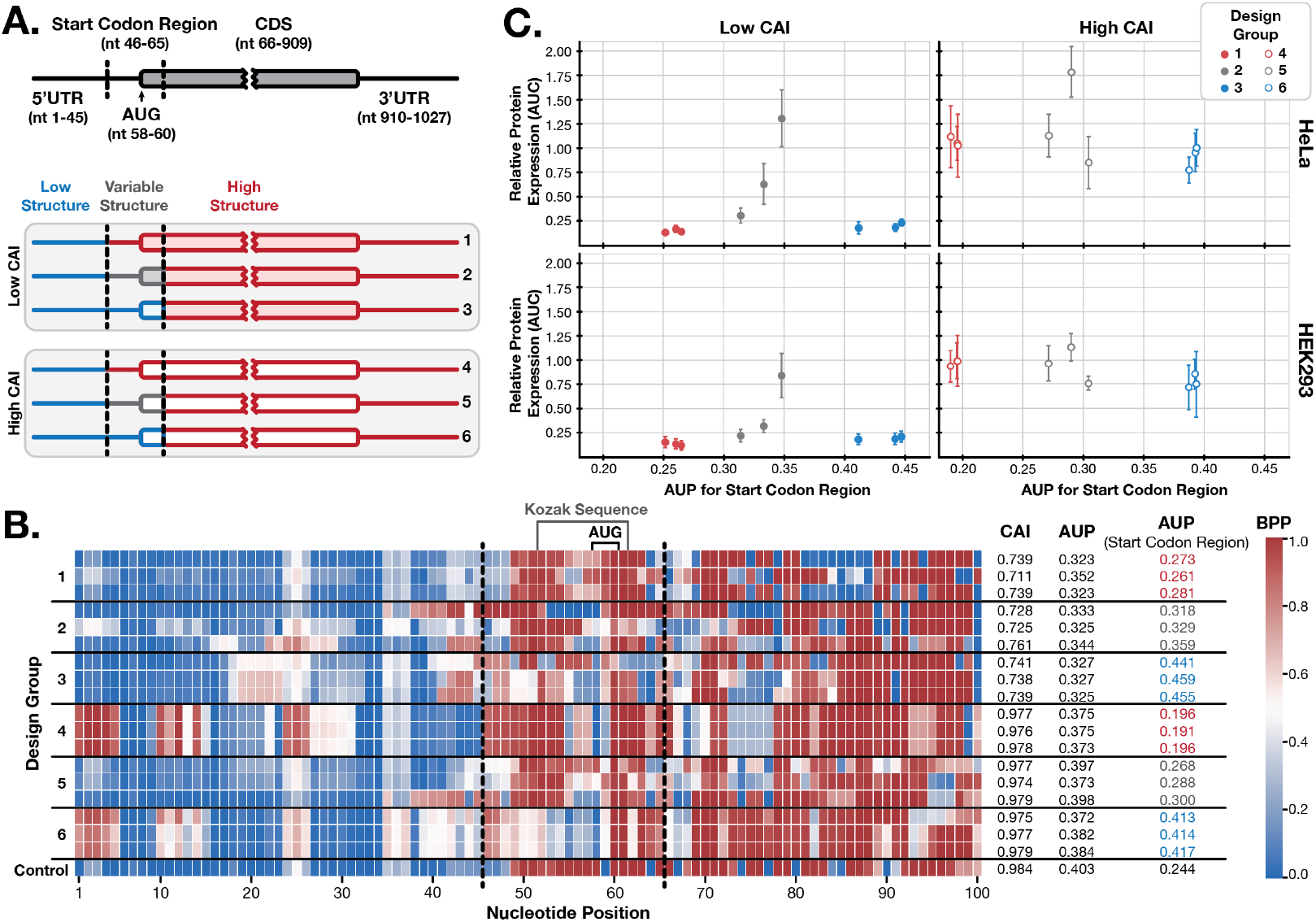
Local structure design. (A) The sequence design scheme for eGFP is depicted for each of the six design groups. (B) A heatmap of base-pairing probability (BPP) per position across the designed sequence shows differences in the predicted structures (zoomed into the first 100 nucleotides for clarity). Less structured regions are blue, while more structured regions are red. CAI and global AUP for the full mRNA are annotated, along with AUP for only the start codon region. (C) Protein expression relative to the control is shown for each sequence, colored by design group.

As expected from suboptimal folding with limited randomness, the base-pairing patterns across the designed sequences were consistent among the three sequence replicates within each design group and generally followed the intended high–medium–low AUP trend (more red, mixed, more blue) within the design window (positions 46-65) (Figure 9B). Across the remainder of the CDS, the CAI values reflect the underlying backbone design of either low or high CAI, with the high CAI group bearing comparable CAI values to the control sequence. The overall AUPs of the sequences were comparable to the control for the high CAI group, and lower than the control for the low CAI group, but broadly similar within each CAI backbone. Because an unstructured 5’UTR is widely considered a key driver of efficient translation initiation, we minimized structure in this region for all constructs, while still maintaining an identical 5’UTR sequence. All groups exhibited predominantly low base-pairing probabilities (blue) in the upstream portion of the 5’UTR (1-45 nts). The calculated AUP values and CAI scores confirm that the mRNA fold algorithm was successful in independently varying RNA structure and codon usage.

The impact of local structure on translation output was assessed in both HEK293 and HeLa cells by measuring GFP fluorescence over time, summarized as the area under the curve (AUC). As expected, the mRNAs in the high CAI set exhibited total protein expression comparable to the control sequence for both cell types, but the mRNAs in the low CAI set exhibited comparably lower protein expression. Notably, in the low CAI groups, a mid-level AUP in the start codon region could partially rescue protein production. In one striking case within the medium AUP group (AUP for start codon region = 0.35), expression levels were comparable to the high-CAI reference mRNA in both cell types, despite a substantially lower CAI (0.74 vs. 0.98) (Figure 9C, top and bottom left, Table 5). We did not observe substantial differences in expression between design groups in the high CAI set, suggesting codon optimization may buffer against the effects of local secondary structure around the start codon.

**Table 5:** Permutation-test *p*-values for coding start region AUP comparisons within CAI groups and cell types.

| Cell Type | CAI Group | Comparison | *p*-value |
| --- | --- | --- | --- |
| HEK293 | Low CAI | Low AUP vs. Med AUP | 0.000019 |
|  |  | Low AUP vs. High AUP | 0.008504 |
|  |  | Med AUP vs. High AUP | 0.000619 |
|  | High CAI | Low AUP vs. Med AUP | 0.785345 |
|  |  | Low AUP vs. High AUP | 0.019595 |
|  |  | Med AUP vs. High AUP | 0.038362 |
| HeLa | Low CAI | Low AUP vs. Med AUP | 0.000002 |
|  |  | Low AUP vs. High AUP | 0.006644 |
|  |  | Med AUP vs. High AUP | 0.000006 |
|  | High CAI | Low AUP vs. Med AUP | 0.229393 |
|  |  | Low AUP vs. High AUP | 0.110230 |
|  |  | Med AUP vs. High AUP | 0.025453 |

Together, these results suggest that the structure surrounding the start codon can meaningfully influence protein production. They also highlight the ability of our algorithm to systematically design local RNA structural variants to test such hypotheses.

## 6 Discussion

We present here a new mRNA co-optimization and folding tool, `mRNAfold`. This algorithm simultaneously optimizes sequences for MFE and CAI and incorporates several novel features, including suboptimal folding, target unstructured regions, incorporation of UTRs, and parallel execution. Benchmarks show that the `mRNAfold` software package is the fastest non-heuristic algorithm available and is even faster with multiple cores available. Further, it is competitive in time usage with heuristic algorithms like LinearDesign. All of this is achieved while being the most feature complete mRNA folding software package currently available.

We additionally present a series of experimental results that test sequences designed using this new algorithm. These results demonstrate the strength and flexibility of the new algorithm, provide further evidence that increasing structure lengthens RNA half-life in solution, and also probe the utility of variable structure near the start codon.

`mRNAfold` builds on existing mRNA folding algorithms [37, 45, 12, 4]. It addresses several of the key shortcomings we identified in [41] for existing methods. The most important is our suboptimal traceback method (see Section 3.2). Suboptimal folding for RNA secondary structures has previously been described as a revolution in RNA secondary structure prediction [19]. The same technique seems similarly useful for mRNA folding, since it enables sampling of many diverse sequences to test, rather than just one at a time (as in most other techniques) or only those on the Pareto optimal frontier (as in DERNA [12]). It should be stressed that suboptimal mRNA folding is not a straightforward adaptation of techniques from RNA secondary structure prediction.

As mentioned in Section 3.2, exhaustive RNA suboptimal folding algorithms are based on Wuchty’s algorithm [44]. These methods enable users to pick a free-energy threshold *t* and then sample all substructures that are within *t* of the minimum free energy. This technique does not lend itself to mRNA folding, since the mRNA folding objective is not free energy but instead a combination of CAI and free energy. As such, it is hard for a user to pick *t* as it is not in interpretable units. Further, `mRNAfold`’s suboptimal folding considers all combinations of both sequence and structure, whereas RNA secondary structure suboptimal folding considers only structure. Consequently, the size of the space grows much faster so it becomes intractable quickly when *t* > 0. This motivated us to develop a different suboptimal paradigm that finds the top-*K* solutions using a priority queue-based strategy.

A popular stochastic sampling method for RNA secondary structures was given by Ding & Lawrence [7] that draws samples from the probability-weighted ensemble of structures. We cannot use this method to generate suboptimal folds for mRNA, since it relies on the partition function, which is not well defined for mRNA folding.

Section 3.3 provides a method to execute mRNA folding in parallel. However, this only includes filling the dynamic programming tables and not suboptimal folding. Our software includes a heuristic where multiple cores are used during suboptimal folding, each running a randomized suboptimal fold with a different seed to increase diversity. However, this method wastes time generating duplicates for low randomness parameter values. A more holistic approach may yield better parallel suboptimal folding.

Despite these advances, there remains room for further improvement. Sparse MFE RNA secondary structure prediction algorithms exploit properties in the recursions to enable significant speedups [43, 11, 5, 1]. Because mRNA folding algorithms use related MFE recursions, it seems possible to adapt sparse folding techniques to mRNA folding, yielding a significant speedup.

At present, mRNAs are co-optimized for MFE and CAI. Extending the optimization framework to include additional cost functions could broaden design objectives. These might include forbidden motifs (e.g., cleavage sites), modulating nucleotide content (often GC or U), avoiding repetitive subsequences (including inverse repeats), and limiting long stems [17]. It’s possible the CAIMFE function could be extended to handle some of these. In principle, any cost function that can be computed as a piecewise sum of codons or “nearest neighbor” RNA secondary structure motifs can be included. For instance, GC-content can be incorporated directly by adding weights to edges that lead to G or C nucleotides.

mRNA folding algorithms consistently improve in-solution stability, as demonstrated in this study and prior work [15, 45]. However, these improvements have largely relied on global metrics such as MFE and AUP. While effective, these metrics may overlook finer-grained, local structural features that also contribute to mRNA degradation and function. For example, a study from the Das lab found that apical loop containing Uridines are more susceptible to hydrolysis than symmetric internal loops [15], suggesting that the types and spatial arrangement of structural elements matter. Future algorithmic development could move beyond global structure optimization toward designing or enriching for specific local motifs to enhance stability.

Our experiments also demonstrate that local structural modifications in specific regions of an mRNA can influence protein production. We tested the impact of modulating structure surrounding the start codon while maintaining a similar degree of overall structure and codon optimality throughout the RNA. We expect that lower base pairing probability in the start codon region leads to lower translation initiation efficiency and more widely spaced ribosomes. Whereas lower translation initiation efficiency would result in fewer protein molecules made per unit time, more widely spaced ribosomes have been associated with longer mRNA half-life and therefore more total protein. The interplay between these contradictory effects may underlie the complex relationship observed between start codon structure and protein output for the low CAI design groups, where mid-level structure produced higher total protein output than low or high structure groups. The increase in protein output for the mid-level design group was statistically significant, but the variability between different sequences in that group was high. This could suggest that in addition to AUP, the specific type of secondary structure could contribute to initiation rate and mRNA half-life. In the high CAI group we did not observe a significant impact of start codon region structure on protein output, suggesting that variation in initiation rate may play less of a role for mRNAs with extremely high optimality.

The experiments presented here highlight the potential of local secondary structure to impact protein production. Here we modulated structure near the start codon, but future experiments can use `mRNAfold` to modulate UTRs, specific CDS regions, or stop codon context of the mRNA and measure impact on initiation, elongation, and/or termination, as well as RNA half-life. One useful application would be to test a proposed strategy for preventing ribosome collisions, which are known to occur during translation elongation and trigger early ribosome dissociation and mRNA degradation [8]. Interspersed structural elements in the CDS have been proposed as a way of modulating ribosome spacing to prevent collisions [22]. Our algorithm enables the design of such structures, offering a tool to test these hypotheses experimentally.

Current folding algorithms largely ignore the UTRs during the optimization phase and thus could potentially miss the impact of UTR-CDS interactions. `mRNAfold` allows optimizations that are UTR-aware. Our experimental design is an example, but there are many avenues for future work. The techniques we describe in Section 3.5 can be extended to penalize any arbitrary set of base pairs or to reward any set of base pairs. So, `mRNAfold` could be used to optimize the CDS in such a way that the native UTR structures are preserved by penalizing all base pairs that are not in the native structures and rewarding those that are. Alternatively, if one wished to optimize for binding sites between the 3’UTR and CDS, then instead the same technique could be used to encourage those pairs.

Together, the advances in `mRNAfold` open the door to a new generation of structure-aware mRNA design strategies that jointly optimize structural features and codon usage to fine-tune translation and stability. The `mRNAfold` tool provides the computational foundation to enable and explore such integrated design approaches.

## Acknowledgments

We would like to thank David Matthews (University of Rochester Medical Center) and Elena Rivas (Harvard University) for their valuable input and advice during the drafting of this manuscript.

