## Supplementary Materials for "mRNAfold: Co-optimization of Global Stability, Local Structure, and Codon Choice via Suboptimal Folding"

### 1 Formal Derivation of the mRNA Folding Problem

We start with some fundamental definitions. These are largely borrowed from our recent review of mRNA folding algorithms [12], which we suggest the reader consult for details beyond what is presented here.

Given an RNA sequence $\pi$ we can define a set $S$ of valid structures. In the Zuker-Stiegler algorithm, we define a valid structure as a properly nested secondary structure.

Formally, let $\pi$ represent an RNA sequence. An RNA is a sequence of nucleotides denoted by 'A', 'U', 'G', and 'C': $\pi \in \{\mathrm{A}, \mathrm{U}, \mathrm{G}, \mathrm{C}\}^\star$. A valid structure $s \in S$ is a set of pairs representing bonds between nucleotides. Only three nucleotide combinations can pair: AU, GC, GU. Note that these can pair in either orientation, e.g., AU and UA are both valid pairs.

A single nucleotide can be in at most one pair in a valid structure: $(i, j) \in s \implies (x, y) \ni s$ such that $(x, y) \neq (i, j)$ and ($x = i$ or $x = j$ or $y = i$ or $y = j$). A valid structure contains no crossing pairs. Two pairs $(i, j), (k, l) \in s$ cross iff $i < k < j < l$ or $k < i < l < j$.

mRNA folding algorithms generally use an energy function $\Delta G(s|\pi)$ that gives the free energy change of a structure $s$ for the sequence $\pi$.

Algorithms that predict RNA structures often predict the structure with the Minimum Free Energy (MFE). Formally, the MFE for a given RNA sequence $\pi$ is:

$$\mathrm{MFE}(\pi) = \min_s \Delta G(s|\pi). \tag{1}$$

Ties for the minimum are broken arbitrarily.

The energy function employed is commonly based on the Nearest Neighbor (NN) model [11, 5, 4, 6]. Rooted in extensive thermodynamic parameters derived from optical melting studies, this modeling approach dates back to the 1970s [9], and continues to undergo refinement and advancement today [15, 6].

mRNA folding algorithms use $\mathrm{MFE}(\pi)$ as a measure of stability [10, 14, 2, 1, 13]. A sequence $\pi$ is considered to be more stable the lower its $\mathrm{MFE}(\pi)$ value is. In addition, modern mRNA folding incorporate Codon Adaptation Index (CAI) [8] as a measure of codon optimality and thus protein expression [14, 2].

Codon Adaptation Index (CAI) measures adaptation of a sequence to the host organism [8]. It assigns every codon a weight:

$$\mathrm{CW}(c) = \frac{f_c}{\max_{s \in \textsc{Synonymous}(c)} [f_s]}.$$

Which is the usage frequency $f_c$ of the codon $c$ divided by the the maximum frequency over the set of all synonymous codons $\textsc{Synonymous}(c)$. Codon usage frequencies are derived from reference mRNA transcript data for a set of highly expressed genes in a target organism. CAI is a geometric mean of codon weights:

$$\mathrm{CAI}(\pi) = \sqrt[|\alpha|]{\prod_{1 \leq i \leq |\alpha|} \mathrm{CW}(c_i)}. \tag{2}$$

CAI values range from 0 to 1 indicating low and high adaptation respectively. In Equation (2), let $\pi$ be a CDS, $\alpha$ be the protein $\pi$ codes for, and $c_i$ be the codon for the $i$-th amino acid. Denote the length (in codons or amino acids) of $\alpha$ by $|\alpha|$. It is convenient to represent CAI in logarithmic form:

$$\log(\mathrm{CAI}(\pi)) = \frac{1}{|\alpha|} \sum_i \log(\mathrm{CW}(c_i)).$$

The $\frac{1}{|\alpha|}$ term is omitted when calculating the combined CAI and MFE objective, which is explained next. That is, we use the unnormalized $\log(\mathrm{CAI})$:

$$\mathrm{LCAI}(\pi) = \sum_i \log(\mathrm{CW}(c_i)). \tag{3}$$

To find the optimal CDS that balances MFE and codon usage, mRNA folding algorithms combine $\log(\text{CAI})$ and MFE into a single objective score:

$$\text{CAIMFE}(\pi) = \text{MFE}(\pi) - \lambda \text{LCAI}(\pi). \tag{4}$$

We drop the $\frac{1}{|\alpha|}$ term from $\log(\text{CAI})$ in CAIMFE since MFE grows linearly with sequence length, so it is natural to scale CAI by $\lambda \times |\alpha|$. Observe that $|\alpha| \times \frac{1}{|\alpha|}$ cancels.

Now we can define a combined MFE and CAI mRNA folding problem:

$$\text{mRNAFold}(\alpha) = \text{argmin}_{\pi \in \text{CDS}(\alpha)} \text{CAIMFE}(\pi). \tag{5}$$

#### 2 Dynamic Programming Complexity Analysis

When designing an mRNA of length $N$, the number of nodes in the codon graph is bounded by $\mathcal{N} = 4N = O(N)$ assuming the standard codon table with 4 nucleotides. Thus, the $P(u, v)$ table is $O(N^2)$ in size since there are $\binom{\mathcal{N}}{2}$ combinations of $u$ and $v$. The same applies to the LCAI($u \rightsquigarrow v$) table, since the table contains all combinations of $u$ and $v$. For $M(c, u, v)$, observe that $c$ only has 3 possible states, $c \in \{0, 1, 2\}$, so the size of the table is also $O(N^2)$ since there are $3 \times \binom{\mathcal{N}}{2}$ combinations of $c, u, v$. For $E(u)$, the size of the table is $O(N)$.

Filling the table for $P(u, v)$ is dominated by looping over all $k, l$ combinations and the associated $n_k, n_l$ nodes. This amounts to all pairs of nodes between $u$ and $v$, which is $O(N^2)$. However, we truncate the number of unpaired nucleotides in a TwoLoop to 30 since large loops have high free energies. This is common practice in RNA folding algorithms [7, 3]. This limits the complexity to $O(1)$, albeit with a large constant factor. So the total cost to compute $P$ is $O(N^2)$ due to the number of $u, v$ combinations.

Filling the table for $M(c, u, v)$ involves looping through split points $k$ and associated nodes $n_k, n_{k+1}$. This amounts to all pairs of adjacent nodes between $u$ and $v$, which is $O(N)$. So the total cost to compute $M$ is $O(N^3)$, since there are $O(N^2)$ combinations of $c, u, v$ and each takes $O(N)$ time to compute.

The analysis for $E(u)$ is similar to $M(c, u, v)$, but we do not need to track $v$ since $E(u)$ computes the answer for a suffix of the graph. There are $O(N)$ options for $u$ and each requires $O(N)$ time to loop over split points, so the total time complexity is $O(N^2)$.

Filling the table $M$ dominates the complexity of the dynamic programming, so the total time complexity is $O(N^3)$ and total memory complexity is $O(N^2)$.

#### 3 Complete Suboptimal Folding Algorithm

##### 3.1 Relevant Information from the Main Text

The pseudocode in Algorithm S1 provides an overview of the suboptimal folding algorithm. The logic is broken up into Trace$E$, Trace$P$, Trace$M$, and TraceLCAI. These functions recapitulate the dynamic programming recursions. The same basic principles are used in each to translate from dynamic programming to traceback.

It is worth pointing out that the LCAI table is included in the traceback. It is easy to miss the significance of this, which is that it allows us to recover the sequence identity of stretches of unpaired nucleotides.

The suboptimal folding pseudocode makes use of $\text{Add}_{(F,S)}(\hat{t}, \delta_F, \delta_S) = \{\hat{\pi}, \hat{s}, F \cup \delta_F, S_{\text{part}} + \delta_S\}$, which adds the states subset $\delta_F$ to $F$ and adds the combined energy and log-CAI score $\delta_S$ to $S_{\text{part}}$. Similarly, the $\text{AddBase}(\hat{\pi}, b_{n_i})$ adds the base $b_{n_i}$ to position $i$ in the partial sequence $\hat{\pi}$, and $\text{AddPair}(\hat{s}, i, j)$ inserts a base pair $(i, j)$ to a partial structure $\hat{s}$.

##### 3.2 Psuedocode

**Supplementary Algorithm S1** Suboptimal traceback algorithm

```
procedure InitTrace(α)
    Input: An amino acid sequence α
    Output: Global state is initialized to do tracebacks
    Fill the dynamic programming tables LCAI(u ⇝ v), E(u), P(u, v), and M(p, u, v)
    Q = {}                                          ▷ Priority queue starts empty
    for u ∈ atpos_1 do                              ▷ Add all initial states to the priority queue
        Push(Q, ⟨∅, ∅, {E(u)}⟩)                     ▷ Empty partial sequence and structure
    end for
end procedure
procedure NextTrace( )
    Input: No input but depends on InitTrace() setting up global state
    Output: The next lowest scoring traceback if there is one or an empty traceback. Modifies the queue
    if Empty(Q) then
        return ∅                                    ▷ Nothing to traceback
    end if
    loop                                            ▷ Keep looping until front of the queue is a complete traceback
        t̂ ← PopMin(Q)                               ▷ Pop the min priority from the queue
        if t̂.x = ∅ then                             ▷ If there are no states left to expand, loop is over
            return ⟨t̂.π̂, t̂.ŝ⟩
        end if
        ε ← PopState(t̂.x)                           ▷ Pop one of the states to expand next
        if ε is from LCAI then
            TraceLCAI(ε, t̂)
        else if ε is from P then
            TraceP(ε, t̂)
        else if ε is from M then
            TraceM(ε, t̂)
        else if ε is from E then
            TraceE(ε, t̂)
        else
            crash, ε is an invalid state
        end if
    end loop
    while |Q| > K do                                ▷ While the queue is too large, remove the maximum score partial traces
        PopMax(Q)
    end while
end procedure
```

**Supplementary Algorithm S2** Traceback for LCAI($n_i \rightsquigarrow n_j$).

**procedure** TRACELCAI($\varepsilon, \hat{t}$)
  **Input:** A state $\varepsilon$ in LCAI, and the partial trace $\hat{t}$ that $\varepsilon$ came from
  **Output:** Modifies the global priority queue, $Q$, by expanding $\varepsilon$
  $n_i, n_j \leftarrow \varepsilon.n_i, \varepsilon.n_j$ ▷ Rename for convenience
  $\hat{t} \leftarrow$ ADDBASE($\hat{t}.\hat{\pi}, b_{n_i}$) ▷ Add the $i$-th base to the sequence
  **if** $i = j$ **then** ▷ Handle base cases for recursion
    **if** $n_i = n_j$ **then**
      PUSH($Q, \hat{t}$) ▷ Add back to queue (minus this base case) without any added states
    **end if**
    **return**
  **end if**
  **for** $n_{i+1} \in \text{out}_{n_i}$ **do** ▷ Look at all possible next nodes
    PUSH($Q$, ADD$_{(F,S)}$($\hat{t}$, {LCAI($n_{i+1} \rightsquigarrow n_j$)}, 0))
  **end for**
**end procedure**

**Supplementary Algorithm S3** Traceback for $P(n_i, n_j)$.

**procedure** TRACE$P$($\varepsilon, \hat{t}$)
  **Input:** A state $\varepsilon$ in $P$, and the partial trace $\hat{t}$ that $\varepsilon$ came from
  **Output:** Modifies the global priority queue, $Q$, by expanding $\varepsilon$
  $n_i, n_j \leftarrow \varepsilon.n_i, \varepsilon.n_j$ ▷ Rename for convenience
  **if** $(b_{n_i}, b_{n_j})$ is not a valid base pairing **then** ▷ Base case for recursion: invalid base pairs
    **return** ▷ Ignore this trace as it is invalid
  **end if**
  $\hat{t} \leftarrow$ ADDPAIR($\hat{t}.\hat{s}, i, j$) ▷ Adds the pair between $n_i$ and $n_j$, since $P$ assumes they are paired
  PUSH($Q$, ADD$_{(F,S)}$($\hat{t}$, {LCAI($n_i \rightsquigarrow n_j$)}, ONELOOP($b_{n_i}, b_{n_j}$))) ▷ ONELOOP case
  **for** $k : i < k < j$ **do** ▷ The TWOLOOP cases
    **for** $l : k < l < j$ **do**
      **for** $n_k \in \text{atpos}_k$ **do**
        **for** $n_l \in \text{atpos}_l$ **do**
          $\delta_S \leftarrow$ TWOLOOP($b_{n_i}, b_{n_j}, b_{n_k}, b_{n_l}$)
          PUSH($Q$, ADD$_{(F,S)}$($\hat{t}$, {$P(n_k, n_l)$, LCAI($n_i \rightsquigarrow n_k$), LCAI($n_l \rightsquigarrow n_j$)}, $\delta_S$))
        **end for**
      **end for**
    **end for**
  **end for**
  **for** $n_{i+1} \in \text{out}_{n_i}$ **do** ▷ The multiloop cases
    **for** $n_{j-1} \in \text{in}_{n_j}$ **do**
      PUSH($Q$, ADD$_{(F,S)}$($\hat{t}$, {$M(2, n_{i+1}, n_{j-1})$, LCAI($n_i \rightsquigarrow n_{i+1}$), LCAI($n_{j-1} \rightsquigarrow n_j$)}, $\text{ML}_{\text{init}} + \text{ML}_{\text{c}}$))
    **end for**
  **end for**
**end procedure**

**Supplementary Algorithm S4** Traceback for $M(c, n_i, n_j)$.

**procedure** $\textsc{Trace}M(\varepsilon, \hat{t})$
  **Input:** A state $\varepsilon$ in $M$, and the partial trace $\hat{t}$ that $\varepsilon$ came from
  **Output:** Modifies the global priority queue, $Q$, by expanding $\varepsilon$
  $c, n_i, n_j \leftarrow \varepsilon.c, \varepsilon.n_i, \varepsilon.n_j$ ▷ Rename for convenience
  **if** $i > j$ **then** ▷ Base cases for recursion
    **if** $c > 0$ **then**
      **return** ▷ Skip invalid base cases
    **else**
      $\textsc{Push}(Q, \hat{t})$ ▷ Adds the trace (minus this base case) back the to queue
      **return**
    **end if**
  **end if**
  **for** $n_{i+1} \in \text{out}(n_i)$ **do** ▷ Add unpaired at $i$ case
    $\textsc{Push}(Q, \textsc{Add}_{(F,S)}(\hat{t}, \{M(c, n_{i+1}, n_j), \text{LCAI}(n_i \rightsquigarrow n_{i+1})\}, \text{ML}_u))$
  **end for**
  **for** $k : i < k \leq j$ **do** ▷ Add pair at $(i, k)$ case
    **for** $n_k \in \text{atpos}(k)$ **do**
      **for** $n_{k+1} \in \text{out}(n_k)$ **do**
        $\textsc{Push}(Q, \textsc{Add}_{(F,S)}(\hat{t}, \{P(n_i, n_k), M(\max(0, c-1), n_{k+1}, n_j), \text{LCAI}(n_k \rightsquigarrow n_{k+1})\}, \text{ML}_c))$
      **end for**
    **end for**
  **end for**
**end procedure**

**Supplementary Algorithm S5** Traceback for $E(n_i)$.

**procedure** $\textsc{Trace}E(\varepsilon, \hat{t})$
  **Input:** A state $\varepsilon$ in $E$, and the partial trace $\hat{t}$ that $\varepsilon$ came from
  **Output:** Modifies the global priority queue, $Q$, by expanding $\varepsilon$
  $n_i \leftarrow \varepsilon.n_i$ ▷ Rename for convenience
  **if** $i = N + 1$ **then** ▷ Base case for recursion: past the end of the codon graph; $n_i = \omega$
    $\textsc{Push}(Q, \hat{t})$ ▷ Adds the trace (minus this base case) back the to queue
    **return**
  **end if**
  **for** $n_{i+1} \in \text{out}_{n_i}$ **do** ▷ First case in recurrence, unpaired nucleotide at $n_i$
    $\textsc{Push}(Q, \textsc{Add}_{(F,S)}(\hat{t}, \{E(n_{i+1}), \text{LCAI}(n_i \rightsquigarrow n_{i+1})\}, 0))$
  **end for**
  **for** $k : i < k \leq N$ **do** ▷ Second case in the recurrence, split on base pair
    **for** $n_k \in \text{atpos}_k$ **do**
      **for** $n_{k+1} \in \text{out}_{n_k}$ **do**
        $\textsc{Push}(Q, \textsc{Add}_{(F,S)}(\hat{t}, \{P(n_i, n_k), E(n_{k+1}), \text{LCAI}(n_k \rightsquigarrow n_{k+1})\}, 0))$
      **end for**
    **end for**
  **end for**
**end procedure**

##### 3.2.1 Suboptimal Folding Complexity

To analyze the time complexity of our suboptimal folding method, let's assume that we are using it to generate the top-$K$ tracebacks for an mRNA sequence of length $N$.

The algorithm will call NEXTTRACE() $K$ times. In the worst case, the loop of NEXTTRACE() will iterate $O(N)$ times.

Observe that a single traceback involves looking at $O(N)$ partial traces. Consider all partial traces that contribute to a traceback. In each step, a single state is removed from $x$ and processed. Each state contributes at least one new nucleotide to the partial sequence $\hat{\pi}$, and there are $N$ nucleotides. So there are at most $O(N)$ partial traces for a given trace. As such, $O(N)$ iterations of the main loop in NEXTTRACE are required to produce a single traceback.

Each iteration involves calling a TRACE function. For states referencing the $P$ recurrence, two-loops dominate, but only require $O(1)$ time, albeit with a big constant. For the $M$ and $E$ tables, splitting into a call to $P$ and calls to $M$ or $E$, respectively, involve iteration over all $O(N)$ split points. This dominates the complexity of these tables. Finally, the recurrence for LCAI is constant. Processing a partial trace therefore generates $O(N)$ new partial traces in the worst case.

Each of these new partial traces need to be added to the queue. This costs $O(N + \log K)$ time. This can be broken down into the time to insert each partial trace in the queue and the time to update the partial trace. The queue operation requires $O(\log K)$ time to add the new partial traceback into the queue. The cost of copying $\hat{t}$ then modifying it to add any new nucleotides, structural motifs, or states, is $O(N)$. This is the cost of the ADDSTATES($\hat{t}$, {new states}) function call.

This means that the overall cost of calling NEXTTRACE() is $O(N^3 + N^2 \log k)$. It involves processing $O(N)$ partial traces, each of which may generate $O(N)$ new partial traces, and processing each requires $O(N + \log k)$ time.

The cost of the final loop that maintains the maximum size of the queue amortizes, since the cost of adding each entry to the queue has already been paid.

The total worst case complexity is thus $O(KN^3 + KN^2 \log K)$.

This seems high, but in practice it is usually much better. This analysis assumes all tracebacks are independent, but often they share many partial traces. Also, often a traceback does not require processing $O(N)$ partial traces. That is, many traces overlap in sequence and structure.

### 4 Experimental Assay

#### 4.1 mRNA Stability Assay and Purity Analysis

mRNA constructs were diluted to 0.1 mg/mL in storage buffer (20 mM Tris, 8% sucrose; Teknova) and dispensed into nine replicate 96-well full-skirt PCR plates using an automated liquid handler. Each plate followed an identical, randomized layout of mRNA constructs. After baseline ($t_0$) measurements were collected, two plates were stored at 25°C, and 40°C temperatures to simulate long-term stability conditions.

At $t_0$ and each subsequent time point, plates were removed from storage and analyzed using Fragment Analyzer Capillary Electrophoresis (FACE). Samples (4 $\mu$L) were transferred into 22 $\mu$L of FA Diluent Marker solution using a Hamilton liquid handler. The mixtures were denatured at 72°C for 2 minutes and cooled to 5°C for 5 minutes before being loaded onto the instrument. Separations were performed using 33 cm capillaries and the standard RNA analysis protocol. Measurements were taken every 3–4 days at 40°C, and weekly at 25°C.

#### 4.2 mRNA Transfection and Live-Cell Expression Assays

HEK293 and HeLa cells were seeded in 96-well, clear-bottom, tissue culture–treated plates. HEK293 cells were transfected at a 25ng dose, while HeLa cells were transfected at a 10ng dose. Cells were incubated overnight at 37°C with 5% $CO_2$ to allow adherence. mRNAs were diluted in nuclease-free water to working concentrations such that 1$\mu$L contained the desired dose per well (e.g., 25ng/$\mu$L for a 25ng/well dose).

Lipofectamine™ 2000 (Thermo Fisher Scientific) was diluted 1:50 in Opti-MEM and incubated for 5 minutes at room temperature. The diluted mRNA and Lipofectamine were then mixed in a 1:1 ratio and incubated for 17 minutes at room temperature to allow complex formation. A volume of 20$\mu$L of these complexes was added to each well (in triplicate), and plates were immediately transferred to a Sartorius Incucyte for live-cell imaging. Imaging was performed every 2 hours for 72 hours.

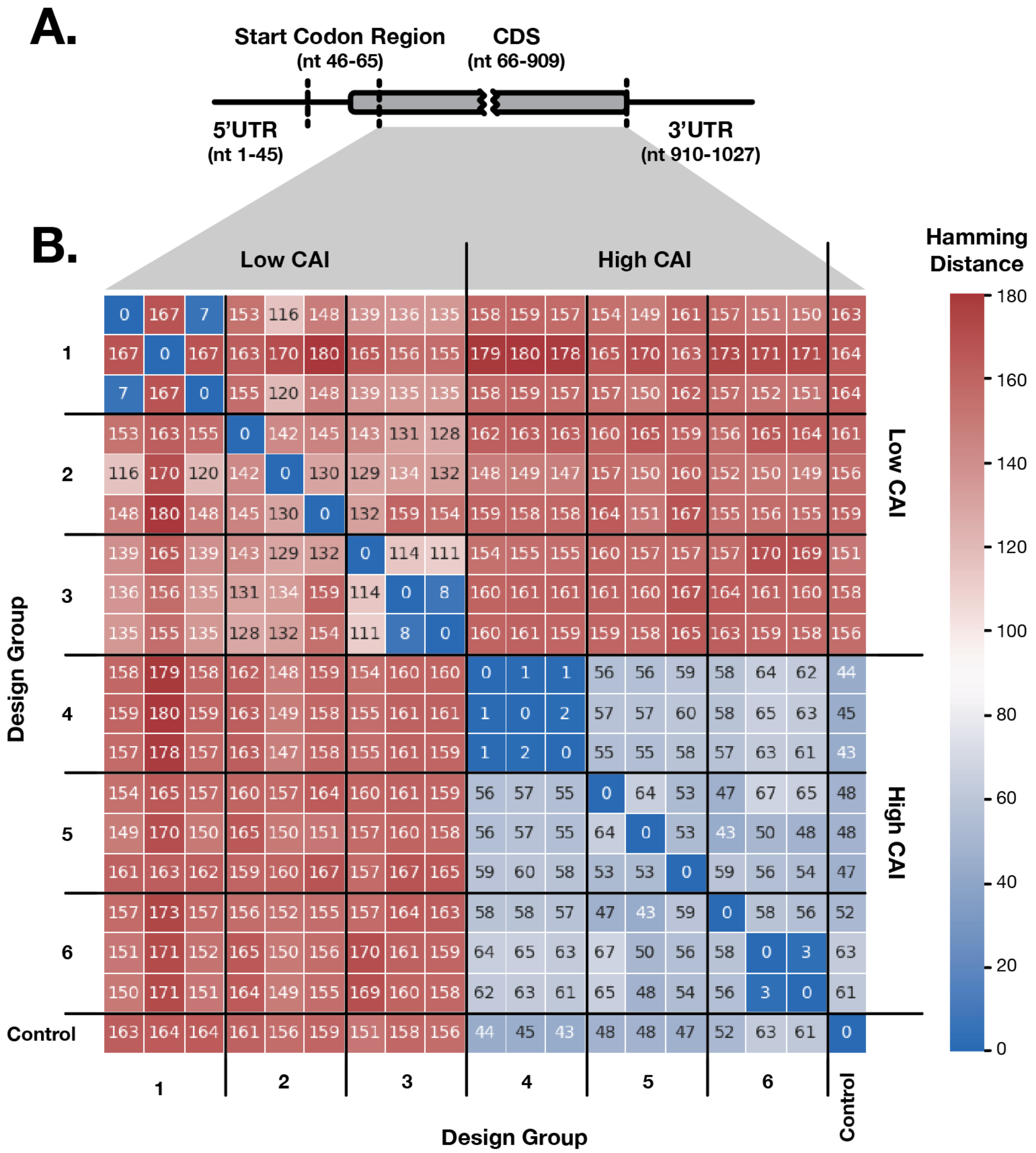


Supplementary Figure S1 Sequence variation in local structure design. (A) We designed eGFP sequence variants targeting low structure in the 5'UTR and variable structure in the start codon region. (B) Since only the coding region was allowed to vary, the designed sequences vary in the remainder of the CDS (not only in the start codon region). The hamming distance between each pair of sequences in the six design groups is shown.

Fluorescence image analysis was conducted using Incucyte software, applying a green-channel surface-area fit model to quantify total green object integrated intensity (TGOII). Expression kinetics were summarized by trapezoidal numerical integration to calculate the area under the curve (AUC) of each fluorescence profile. A GFP-22 mRNA was included on every plate as an internal normalization control, and all expression values were normalized to this reference.
